# TUSC3 serves as a rate-limiting gatekeeper of a glycan-mediated ER Triage Checkpoint for BMP4/Dpp

**DOI:** 10.1101/2025.10.03.680295

**Authors:** Antonio Galeone, Emilio Solazzo, Francesco Lavezzari, Seung-Yeop Han, Bruna My, Riccardo Rizzo, Giuseppe Gigli, Hamed Jafar-Nejad, Thomas Vaccari

## Abstract

Trimming of the three glucose residues decorating nascent *N*-glycoproteins is a critical step for their entry into the endoplasmic reticulum quality control (ERQC) cycle and recognition by ER chaperones. However, the functional relevance of the second glucose (G2) and the regulatory step upstream of its removal by ER glucosidase II (GCS2) remains poorly understood. Here, we report that TUSC3, a component of the oligosaccharyltransferase (OST) complex, regulates G2 to G1 trimming on *N-*glycosylated bone morphogenetic protein 4 (BMP4) and its *Drosophila* homolog Dpp to promote their ERQC entry. Loss- and gain-of-function genetic experiments and biochemical assays in mammalian cells and flies indicate that TUSC3 serves as a dosage-sensitive gatekeeper that influences the decision between proper folding and secretion versus elimination by ER-associated degradation for BMP4 molecules, thereby tuning BMP signaling. Together, these data reveal an unrecognized role for an OST component in early glycoprotein maturation, relevant to a major developmental signaling pathway.

## INTRODUCTION

*N-*glycosylation is a highly conserved co- and post-translational modification that involves the attachment of an oligosaccharide to an asparagine (*N*) residue of a protein via complex biochemical pathways.^1–3^ The vast majority of secretory proteins are *N-*glycosylated in the endoplasmic reticulum (ER) before trafficking to their destinations.^3,4^ In the ER, protein maturation is temporally and spatially organized, and *N-*linked glycans serve as quality control tags that support protein stability, cellular trafficking, and cell-surface interactions.^5–7^ Clients are *N-*glycosylated in the ER by the action of the multimeric oligosaccharyltransferase (OST). The OST complex catalyzes the transfer of Glucose3Mannose9N-acetylglucosamine2 (G3M9GlcNAc2) molecules from dolichol, its lipid carrier, to the consensus sequence N-X-T/S/C (X ≠ P) of the nascent protein. The glucose molecules on the a-antenna of the dolichol-linked oligosaccharides act as a recognition signal to engage the OST complex ^8^ After the attachment, G3M9GlcNAc2-*N-*linked molecules are extensively remodeled by a series of glycosidases and transferases along the secretory pathway.^9^ *N-*glycans’ maturation starts with the removal of the outermost (α1-2)-linked glucose residue by the type II membrane protein glucosidase I (GCS1; MOGS in humans). Glucosidase II (GCS2) then removes the middle (α1-3)-linked glucose residue from G2M9GlcNAc2. This process produces monoglucosylated species (G1M9GlcNAc2) recognized by the lectin chaperones calnexin (CNX) for entering the ER quality control (ERQC).^3,10–12^

In higher eukaryotes, the OST complex is a hetero-oligomer composed of seven highly conserved subunits and exists in two forms, OST-A and OST-B, defined by their respective catalytic subunit STT3A and STT3B.^8,13–15^ The two forms of OST complex share six core subunits, while the seventh subunit is complex-specific: DC2/OSTC or KCP2 for OST-A; TUSC3 or MAGT1 for OST-B.^8,13,14,16^ OST-A primarily mediates co-translational *N*-glycosylation of nascent proteins at the translocon, whereas OST-B catalyzes post-translational *N-*glycosylation of sites skipped by OST-A.^17^

The TUSC3 and MAGT1 subunits possess *N-*terminal regions within the ER lumen with a predicted thioredoxin domain and are both anchored to the ER via *C-*terminal transmembrane domains.^18^ Mass spectrometric analysis of cells and plasma obtained from patients with XMEN disease, which is an X-linked immunodeficiency caused by *MAGT1* mutations,^19^ indicates a selective reduction in *N*-glycan occupancy at specific glycosylation sites.^20^ In parallel, X-ray structural studies of TUSC3 reveal a defined peptide-binding groove adjacent to its active-site cysteine pair that can accommodate peptides in opposite orientations, suggesting a role in modulating the glycosylation efficiency of certain glycosylation sites through interactions with polypeptides.^15,18,21^ In addition, prior work has established that TUSC3 supports STT3B-dependent, site-specific *N-*glycan occupancy, often in cysteine-coordinated/thioredoxin-sensitive contexts.^21,22^ However, it is not known whether TUSC3 also regulates post-transfer *N*-glycan remodeling or otherwise impacts proteostasis.

Genetic and biochemical studies have linked *TUSC3* to tumor suppression in prostate, ovarian, and pancreatic cancers, as well as to the modulation of the unfolded protein response (UPR).^23–27^ In tumor cells, loss of *TUSC3* appears to alter ER stress signaling and chaperone induction and might render tumor cells tolerant to high levels of misfolded glycoproteins.^23,26,28^ *TUSC3* and *MAGT1* have been implicated in Mg²⁺ transport, with possible relevance to *N*-glycosylation.^29,30^ Mutations in *TUSC3* cause an autosomal-recessive congenital disorder of glycosylation (TUSC3-CDG) characterized by intellectual disability, highlighting its physiological importance^13,31,32^ In *Drosophila*, a single homolog of *TUSC3* and *MAGT1* is present, which is called *Ostγ*. A previous study, which referred to this gene as *MagT1*, reported embryonic cuticle defects and adult wing phenotypes in animals with *Ostγ* mutant clones and *Ostγ* knockdown, and proposed potential links to Wingless and Dpp signaling.^33^ However, whether the observed wing phenotypes were indeed caused by impaired Dpp signaling or connected to *N-*glycosylation, protein folding or trafficking was not reported.^33^ All in all, how TUSC3 regulates the interplay between nascent glycans, OST, lectin-chaperones, and ER stress signaling remains poorly understood.

Using mouse cell-based and *Drosophila melanogaster* models, here we provide mechanistic evidence that TUSC3 participates in a triage checkpoint activated by G2M9GlcNAc2 molecules upstream of the ERQC, serving as a bottleneck for selecting proteins eligible to traffic to the Golgi apparatus. To explore the physiological relevance of this triage mechanism, we focused on bone morphogenetic protein 4 (BMP4) and its fly homolog Decapentaplegic (Dpp)— secreted ligands whose maturation and signaling are tightly regulated by ER glycoprotein processing.^34–36^ We found that *TUSC3* and its fly homolog *Ostγ* promote productive engagement of ER chaperones, thereby limiting the accumulation of misfolded/aggregation-prone glycosylated species and supporting ER proteostasis, ultimately ensuring correct levels of BMP4 signaling. Moreover, genetic experiments in flies suggest that TUSC3 regulates additional glycoprotein clients beyond BMP4/Dpp. Our studies provide key insights into the importance of TUSC3 and sugar moieties in mediating a triage checkpoint at an early stage during *N*-glycosylation and in regulating glycoprotein folding and intercellular signaling.

## RESULTS

### *Tusc3* promotes BMP4 signal-sending in mouse embryonic fibroblasts by ensuring proper *N-*glycosylation

BMP4 and its *Drosophila* ortholog Dpp are paradigmatic *N-*glycosylated secreted ligands whose maturation and signaling depend on proper ER processing and a functional ERAD machinery.^34,37–40^ We and others have shown that BMP4/Dpp glycosylation is critical for productive folding, ER exit, and downstream SMAD/Mad activation.^35,36,41^ To test the contribution of TUSC3 to the maturation of *N-*glycosylated cargo, we used a small interfering RNA (siRNA) to knockdown *Tusc3* (the mouse homolog of human *TUSC3*) in mouse embryonic fibroblasts (MEFs), which led to ∼90% reduction in TUSC3 protein level in these cells (Figure S1). We transfected the cells with a double-tagged version of bone morphogenetic protein 4 (*BMP4-HA-Myc*) with an HA tag in its prodomains and a Myc tag in its active domain, allowing us to trace the full-length and active domain of this *N-*glycosylated cargo intracellularly through the secretory pathway and in the culture media after secretion. Upon *Tusc3* knockdown, the conditioned media showed a remarkable decrease in the level of secreted BMP4-Myc active domain compared to control MEFs (Figure 1A). As expected from reduced secretion of BMP4-Myc, *Tusc3*-knockdown MEFs showed impairment in BMP signaling as evidenced by a decrease in the expression of pSMAD1/5 (Figure 1A), which is a BMP effector.^42^ These data indicate that TUSC3 is required for the secretion of the active domain of BMP4 and BMP4 signaling in MEFs.

**Figure 1.**
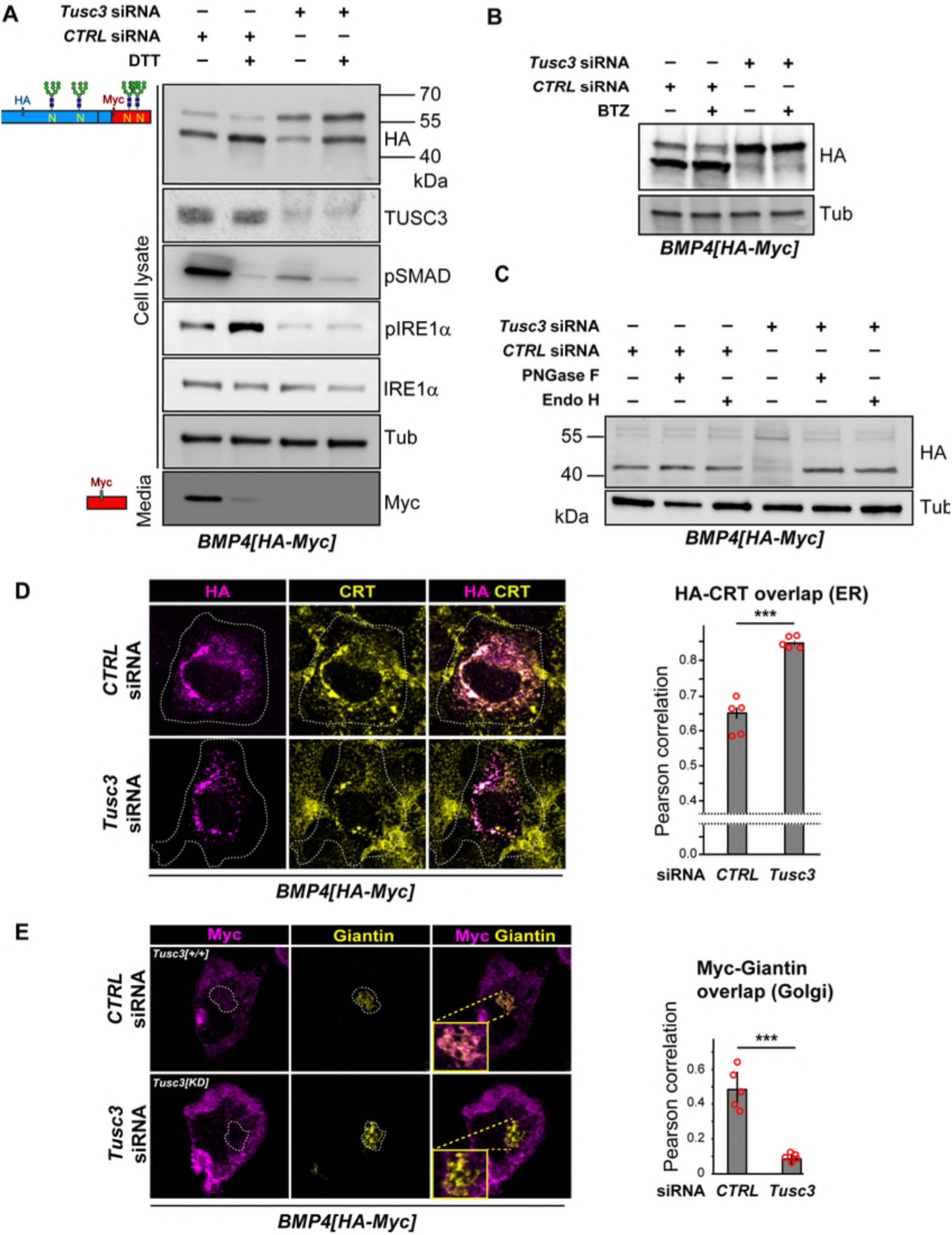
*Tusc3* knockdown impairs *N-*glycosylation–dependent secretion of BMP4 and attenuates ER stress signaling in MEFs. (A) Immunoblot analysis of the indicated markers in MEFs transfected with BMP4-HA-Myc, transfected with control or *Tusc3* siRNA, with or without DTT incubation. Conditioned media were immunoblotted for Myc to detect the secreted BMP4 mature domain. β-Tubulin serves as a loading control. (B) Immunoblot analysis of the indicated markers in MEFs transfected with BMP4-HA-Myc, transfected with control or *Tusc3* siRNA, with or without BTZ incubation. β-Tubulin serves as a loading control. (C) MEFs expressing *BMP4-HA-Myc* and transfected with control or *Tusc3* siRNA were lysed and split into three aliquots: untreated, digested with PNGase F, or digested with Endo H for 1 h at 37 °C. Immunoblotting for HA was used to detected BMP4-HA-Myc. (D) Representative confocal immunofluorescent images of MEFs expressing BMP4-HA-Myc, transfected with control or *Tusc3* siRNA, and stained for HA (green) to detect BMP4 and calreticulin (CRT; red) as an ER marker. Right: quantification of Pearson’s correlation coefficient for HA/CRT co-localization (mean ± SD, n = 5 cells from three independent experiments; ***p < 0.001, Student’s t test). (E) Representative confocal immunofluorescent images of MEFs expressing BMP4-HA-Myc, transfected with control or *Tusc3* siRNA, and stained for Myc (magenta) and the Golgi marker Giantin (yellow). Representative confocal micrographs (63×; scale bar = 10 µm) are shown. Right: quantification of Pearson’s correlation coefficient for Myc/Giantin co-localization (mean ± SD, n = 5 cells pooled from three independent experiments; ***p < 0.001, Student’s t test).

Human BMP4 and its homologs in other animals have four *N*-glycosylation sites, which have been experimentally verified in some cases.^43^ The full-length BMP proteins are glycosylated in the ER and are cleaved by protein-convertase in the Golgi apparatus.^44^ In agreement with our previous report,^41^ the full-length BMP4 protein from MEFs migrates as two bands in immunoblots: a weaker upper band consisting of *N-*glycosylated molecules and a stronger lower band composed of non-glycosylated and/or de-*N*-glycosylated molecules (Figure 1A). Upon *Tusc3* knockdown, HA immunoblotting showed the accumulation of the upper band, compatible with retention of the *N*-glycosylated form of the full-length BMP4-HA-Myc. We previously reported that overexpression of *BMP4-HA-Myc* in control MEFs leads to the accumulation of misfolded BMP4 molecules, upregulation of phosphorylated IRE1α (pIRE1α)—an indicator of unfolded protein response activation,^45,46^—and induction of OS9 and BiP, chaperone-assisting molecules that are crucial for ER associated degradation (ERAD).^41,47–49^. Treating control MEFs with dithiothreitol (DTT) led to a strong induction of IRE1α phosphorylation without an obvious change in the total IRE1α level (Figure 1A), in agreement with induction of ER protein misfolding by DTT.^49^ Surprisingly, *Tusc3*-knockdown MEFs showed a strong reduction in pIRE1α levels compared to control MEFs in both basal and DTT-treated conditions, suggesting insensitivity to ER stressors. Loss of pIRE1α induction in *Tusc3*-knockdown MEFs prompted us to examine whether the BMP4 molecules in *Tusc3*-knockdown MEFs were accumulated due to impairment of ERAD. As shown in Figure 1B, the level of BMP-HA-Myc upper band in *Tusc3*-knockdown MEFs did not further increase upon proteasomal inhibition, suggesting that the BMP4 molecules that accumulate in *Tusc3*-knockdown MEFs were not able to undergo ERAD.

The slow migration of the accumulated BMP4 molecules in immunoblots from *Tusc3*-knockdown MEFs and the lack of pIRE1α induction suggested that these cells accumulate a glycosylated form of BMP4 in the ER. To test this notion, we digested the cell lysates with PNGase F and Endo H enzymes.^50^ Digestion with these enzymes did not alter the migration pattern of the BMP-HA-Myc lower band in control MEFs (Figure 1C), confirming that the molecules present in the lower band lack *N*-glycans.^41^ Importantly, upon digestion of the *Tusc3-*knockdown cell lysates with either enzyme, the BMP4-HA-Myc upper band migrated similarly to the lower band of control MEFs (Figure 1C), indicating that BMP4 retains ER-type (high mannose) *N*-glycans in *Tusc3*-knockdown MEFs. Double staining for HA and the ER marker Calreticulin (CRT)^51^ showed a partial localization of BMP4-HA-Myc to the ER in control MEFs (Figure 1D). This is expected, as at each point in time some BMP4 molecules will be in the ER while others have continued their exocytic trafficking to the Golgi apparatus and beyond. In contrast, BMP4-HA-Myc mostly colocalized with CRT in *Tusc3*-knockdown MEFs (Figure 1D). In addition, double staining for Myc and the Golgi marker Giantin^52^ revealed a limited colocalization that might represent molecules in transit towards secretion. This colocalization is reduced in *Tusc3*-knockdown MEFs compared to control MEFs (Figure 1E). Together, these biochemical and imaging data indicate that *Tusc3* knockdown leads to the entrapment of glycosylated BMP4 molecules in the ER and impairs the activation of pIRE1α, suggesting that TUSC3 plays a role in the activation of a branch of the UPR involved in ER stress response.

### TUSC3 promotes ER-Glucosidase 2 (GCS2)-mediated trimming of G2M9GlcNAc2-BMP4

Our observation of glycosylated-BMP4 accumulation in *Tusc3*-deficient MEFs without activating the ER stress response prompted us to examine whether the glucose residues have been properly trimmed from BMP4 *N*-glycans, considering that *N*-glycoproteins enter the quality control calnexin/calreticulin cycle upon removal of the two distal glucose residues from *N*-glycans. To this end, we used two independent strategies to further enrich BMP4 molecules in the ER by slowing the protein trafficking from ER to the Golgi: (1) culturing the cells at 20 °C, and (2) incubating the cells with the fungal metabolite Brefeldin A, a reversible inhibitor of intracellular vesicle formation.^53^ Moreover, MEFs were treated with NBDNJ,^54^ an inhibitor of ER α-glucosidases I and II (GCS1 and GCS2, respectively) as a positive control.

As shown in Figure 2A, cell extracts from control MEFs cultured at 20 °C displayed five BMP4-HA bands with different degrees of intensity. The lowest one corresponds in size to unglycosylated/de-*N*-glycosylated BMP4 (see Figure 1C). This band slightly increased in intensity after DTT treatment, suggesting that it is likely composed of the deglycosylated form of misfolded BMP4 molecules resulting from ERAD retrotranslocation (Figure 2A, red arrow). Notably, upon GCS inhibition, the intensity of the top three bands increased and the intensity of the fourth band from the top decreased (Figure 2A). Since inhibition of GCS1 and GCS2 inhibits glycose trimming from *N*-glycans, the data suggest that the remaining bands are G3M9GlcNAc2-BMP4 (G3M9), G2M9GlcNAc2-BMP4 (G2M9), G1M9GlcNAc2-BMP4 (G1M9) and M9GlcNAc2-BMP4 (M9)-BMP4 molecules (shared cartoon Figure 2A and B). Rather surprisingly, upon *Tusc3* knockdown, the BMP4-HA-Myc band profile appeared very similar to that resulting from GCS inhibition (Figure 2A), suggesting that *Tusc3*-deficient MEFs accumulated glucosylated M9GlcNAc2-BMP4 molecules in the ER. DTT treatment increased OS9 (Figure 2A), a lectin upregulated during ER stress that recognizes misfolded glycoproteins and delivers them to the retrotranslocon.^32^ In GCS-inhibited MEFs, OS9 levels were less than that observed in DTT-treated MEFs and looked comparable to control cells. However, the level of OS9 in *Tusc3* knockdown cells was less than that in the control MEFs (Figure 2A). These results indicate a functional dissociation between glucosylated-BMP4 buildup in *Tusc3* knockdown and canonical ERAD surveillance: upon *Tusc3* knockdown, the cells fail to sense *N-*glycoprotein misfolding, indicating that terminal glucose residues on the M9GlcNAc2 glycoform of BMP4 need to be removed to activate ERAD-mediated clearance of misfolded BMP4.

**Figure 2.**
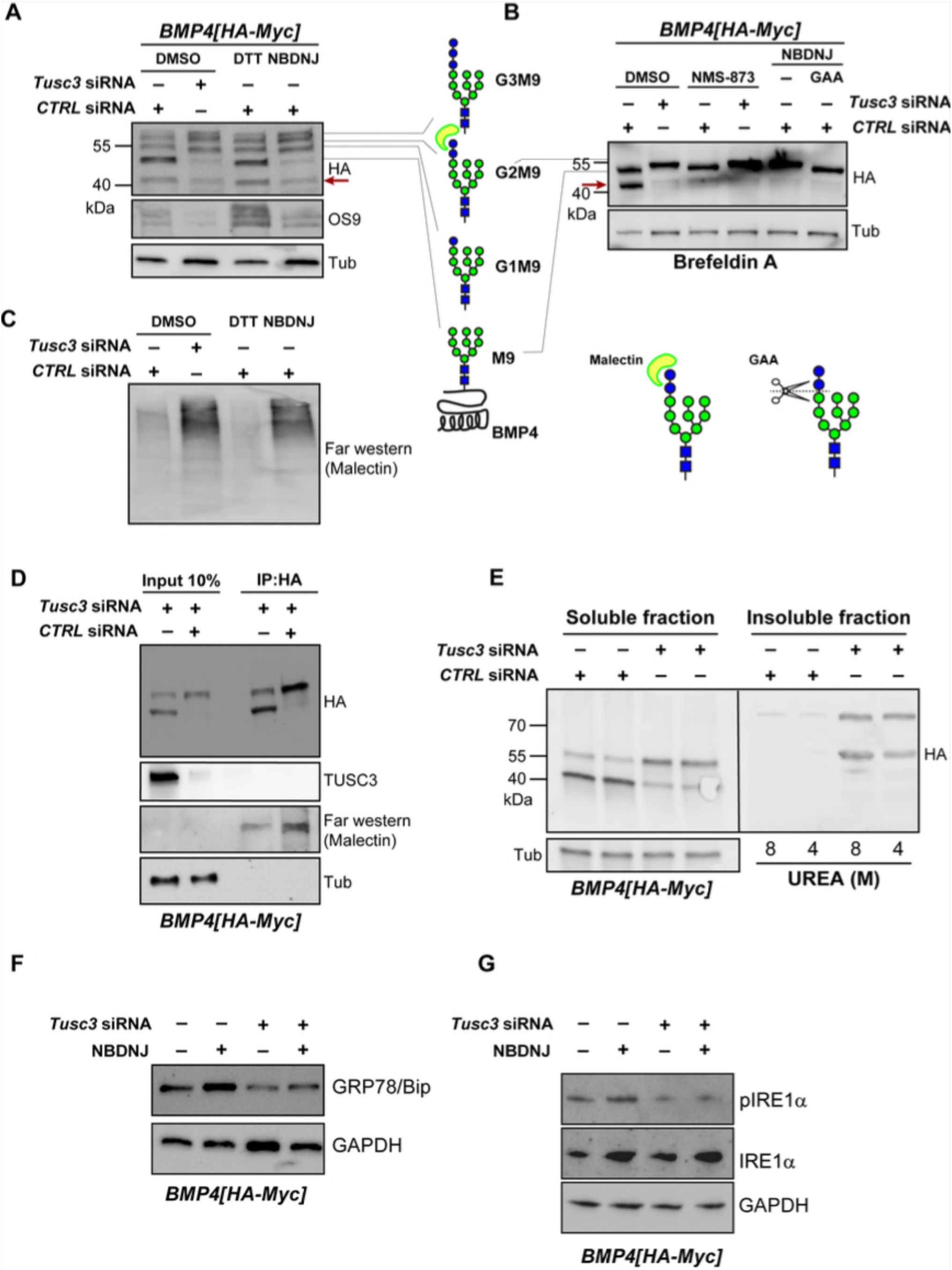
TUSC3 promotes GCS2-mediated glucose trimming and prevents insoluble G2M9-BMP4 accumulation. (A) Western blot with anti-HA in MEFs cultured at 20 °C and treated with DTT for 6 h (to induce misfolding/ERAD), 2 mM NBDNJ for 16 h (inhibitor of GCS1/GCS2), or DMSO as control. The five discrete BMP4-HA species corresponding (top to bottom) to G3M9GlcNAc2-BMP4, G2M9GlcNAc2-BMP4, G1M9GlcNAc2-BMP4, M9GlcNAc2-BMP4, and non-glycosylated/deglycosylated BMP4 (red arrow) are marked. Tubulin serves as loading control. Representative of two independent experiments. (B) HA immunoblots of lysates from MEFs transfected with CTRL or *Tusc3* siRNA and treated with the indicated compounds are shown. GCS inhibitor NBDNJ, 8 h; acid α-glucosidase (GAA), 50 mU/mL, 1 h at 37 °C; VCP inhibitor NMS-873, 10 µM, 6 h. Representative of two independent experiments. (C) Malectin far-western blot of total protein lysates from MEFs under the indicated conditions. (D) HA immunoprecipitation of BMP4-HA from control and *Tusc3*-knockdown MEFs, followed by malectin far western blot. Representative of two independent experiments. (E) Western blot analysis of BMP4-HA in soluble and insoluble protein fractions from control and *Tusc3*-knockdown MEFs. Representative of two independent experiments. (F-G) Western blot of BiP and pIRE1α in MEFs treated with NBDNJ.

When ER-to-Golgi trafficking was impaired by Brefeldin A, BMP4-HA bands migrated slower in both *Tusc3* knockdown and GCS2-inhibited MEFs (Figure 2B). We next digested protein extracts from NBDNJ-treated MEFs with acid α-glucosidase (GAA), a pan-glucosidase expected to remove glucose molecules from glycoforms. Upon GAA treatment, migration of BMP4-HA band was downshifted to match that of the upper band in control cells, confirming the presence of glucose molecules on BMP4-HA-Myc *N*-glycans in extracts of MEFs treated with NBDNJ and likely those depleted of *Tusc3* (Figure 2B). Upon inhibition of VCP, a key component of ERAD,^55,56^ MEFs did not display the lower BMP4-HA band when compared to non-treated control MEFs, providing further evidence that BMP-HA molecules are retrotranslocated in ERAD-dependent manner. Notably, in *Tusc3*-deficient MEFs, the migration of BMP-HA did not change upon VCP inhibition when compared to non-treated *Tusc3*-deficient MEFs (Figure 2B, Supplementary Figure S2A). Together, these observations further suggest that *Tusc3* knockdown leads to the accumulation of BMP4 molecules with glucose-bearing *N*-glycans that fail to be targeted for ERAD.

Given the similarities between the BMP4-HA migration patterns upon *Tusc3* knockdown and GCS inhibition, we next asked whether *Tusc3* knockdown affects malectin binding of *N*-glycoproteins in the corresponding cell lysates. Malectin is a carbohydrate-binding protein that is highly specific for G2M9 glycans and binds glycoproteins before their entry into ERQC.^57^ As expected, treating MEFs with the GCS inhibitor NBDNJ led to a dramatic increase in malectin binding of the cell extracts (Figure 2C). DTT treatment did not lead to increased malectin binding compared to control cells (Figure 2A), indicating that despite the significant protein misfolding expected to arise from DTT treatment, GCS2 is able to efficiently remove the second glucose residue from *N*-glycans decorating misfolded *N*-glycoproteins. In contrast, malectin exhibited broad and strong binding to *N*-glycoproteins in the total protein extract of MEFs depleted of *Tusc3*, similar to the MEFs subjected to GCS inhibition (Figure 2C). These observations strongly suggest that BMP4 molecules harboring G2M9 *N*-glycans accumulate in *Tusc3*-deficient cells. The data also suggest that NBDNJ-treated cells still have some GCS1 activity but show a strong inhibition of GCS2, hence the accumulation of G2M9 glycans.

Seeking further evidence of G2M9-BMP4 accumulation upon *Tusc3* knockdown, we used HA immunoprecipitation to enrich HA-tagged BMP4 molecules and tested the binding of BMP4 to malectin by using the far western technique.^58^ Control MEFs showed malectin-positive glycosylated BMP4, suggesting that some BMP4-HA molecules harbor G2M9 *N*-glycans in these cells (Figure 2D). The immunoprecipitated BMP4-HA from *Tusc3*-deficient MEFs showed a higher level of malectin binding, parallel to the higher levels of BMP4-HA in these cells (Figure 2D). Together, these observations provide compelling evidence that *Tusc3*-deficient MEFs accumulate BMP4-HA molecules at a stage before GCS2-mediated removal of the second glucose residue.

Compared to monomeric misfolded proteins, which still retain the ability to be folded by chaperones or be degraded, insoluble proteins are more prone to aggregation.^59^ Given the observations that *Tusc3*-deficient MEFs did not induce ER chaperones (Figure 1A and Figure 2A) and accumulated G2M9-BMP4, we asked whether G2M9-BMP4 molecules tend to form aggregates of insoluble molecules. As shown in Figure 2E, analysis of soluble and insoluble fractions of crude protein extracts showed a remarkable increase of BMP4 molecules in the insoluble fraction from *Tusc3* knockdown MEFs compared to control MEFs. To further assess whether accumulation of insoluble BMP4 molecules in *Tusc3*-deficient cells triggers canonical ER stress responses, we measured the levels of BiP and pIRE1α. Treatment of MEFs with NBDNJ resulted in a strong upregulation of both BiP and pIRE1α (Figure 2F and G, Supplementary Figure S2B), indicating the activation of the UPR due to the accumulation of unfolded glycoproteins. By contrast, *Tusc3*-knockdown cells exhibited significant reduction of both BiP and pIRE1α levels (Figure 2F and G, Supplementary Figure S2B), thereby suggesting that upon *Tusc3* knockdown, insoluble G2M9-decorated proteins fail to engage ER chaperone surveillance mechanisms, potentially due to their aggregation into chaperone-inaccessible forms. Importantly, these findings also reveal a functional distinction between G2M9 accumulation caused by glycosidase inhibition and by *Tusc3* knockdown. Although both conditions result in comparable buildup of G2M9-decorated glycoproteins, the NBDNJ-treated cells robustly activate the UPR via BiP and pIRE1α, whereas Tusc3-deficient cells fail to do so.

### Fly *Ostγ* has a high level of functional conservation with human TUSC3 in mediating BMP signaling

Since the above observations were based on overexpression of *BMP4-HA-Myc* in *Tusc3*-deficient MEFs, we asked whether *TUSC3* physiologically regulates endogenous BMP signaling in an *in vivo* context, and if so, whether it does so by a similar mechanism to that uncovered in MEFs. To address this point, we used transposon-mediated mutagenesis to generate two imprecise excision alleles of the *Drosophila* Oligosaccharide transferase γ subunit (*Ostγ*, Figure 3A), which shares a high degree of protein sequence identity with human *TUSC3*. Genetic and molecular characterization by PCR indicated that the isolated alleles, *OstγΔ4* and *OstγΔ5*, are severe hypomorphic or null alleles and show 100% lethality in homozygosity as well as compound heterozygosity with each other or a Deficiency strain uncovering *Ostγ* (Figure 3A and 3B, Figure S3).

**Figure 3.**
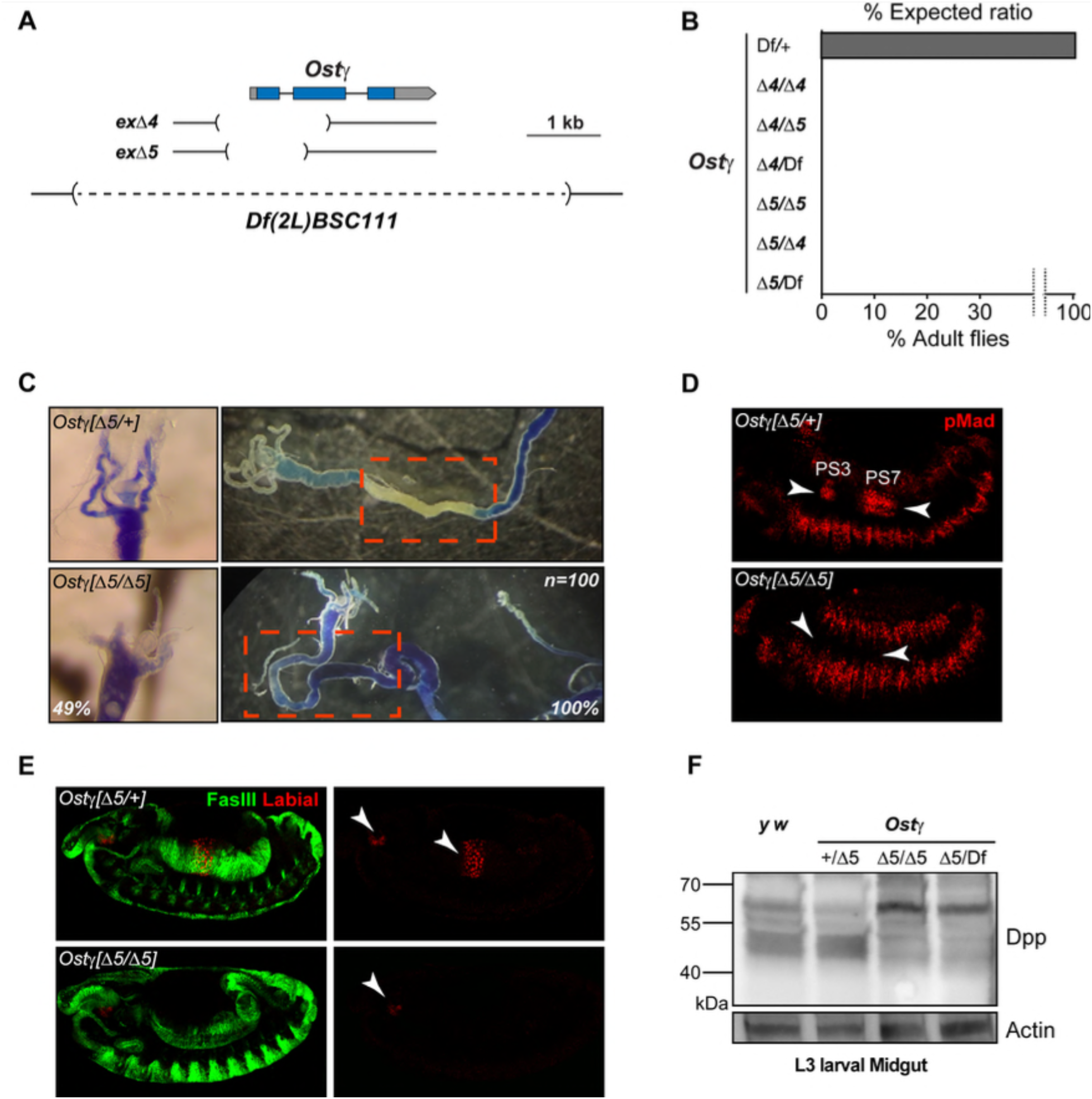
Loss of fly *Ostγ* impairs Dpp/BMP signaling in the embryonic visceral mesoderm and larval midgut. (A) Schematic of the *Ostγ* locus and imprecise excision alleles (*Δ4* and *Δ5*). Exons are represented as boxes, with introns depicted as lines. (B) Survival assay showing percentage of eclosed adults from balanced heterozygous progeny (*OstγΔ/+ × OstγΔ/+*). n ≥ 200 embryos scored per genotype. (C) Third-instar larval midguts from *OstγΔ5/+* (control) or *OstγΔ5/Δ5* were dissected and fed bromophenol blue (BPB)– supplemented food for 2 h. BPB turns yellow in the acidic region (acid zone, AZ). Note the distinct yellow AZ (dashed box) in control midguts. n = 100. (D) Stage-14 embryos (ventral view) from *OstγΔ5/+ and OstγΔ5/Δ5* were fixed and stained with anti–pMad (red). PS3 and PS7 are marked with white arrowheads. (E) Stage-14 embryos from *OstγΔ5/+* and *OstγΔ5/Δ5* were stained for Labial (red) and Fas3 (green) to mark VM boundaries. White arrowheads mark Labial expression in PS3 and PS7. (F) Western blot of total protein lysates from third-instar larval midguts (genotypes: *yw, OstγΔ5/+, OstγΔ5/Δ5*, and *OstγΔ5/Df(2L)Exel7030*) probed with an anti-Dpp antibody recognizing the prodomain. Tubulin serves as a loading control. Representative of two independent biological replicates.

Among the transforming growth factor beta (TGF-β) superfamily, decapentaplegic (dpp) is the *Drosophila* ortholog of BMP2/4.^60^ It has been reported that Dpp and BMP4 share several developmental functions and conserved roles in establishing and patterning the dorsal-ventral axis of several organs.^61^ In fly embryos, Dpp is expressed in narrow bands in parasegments 3 (PS3) and PS7 of the visceral mesoderm (VM). From there, it activates BMP signaling in the neighboring endoderm and induces the homeodomain gene *labial* in PS7, which is responsible for forming the acid zone region of the midgut.^34,62,63^ To test whether BMP signaling is impaired in the larval midgut of *Ostγ* mutants, we carried out bromophenol blue (BPB) feeding assays on *Ostγ ^Δ5/Δ5^* and compared it with *Ostγ ^Δ5/+^* siblings as a control. BPB is a nontoxic compound that can be mixed with fly food and turns yellow from blue as the pH is decreased from 7.0 to 1.0. Compared to control, *Ostγ ^Δ5/Δ5^* larvae showed a fully penetrant loss of yellow color in the midgut, indicating the loss of the acid zone (Figure 3C). Moreover, around half of the mutant larvae exhibited severe shortening of the gastric caeca (Figure 3C), another part of the larval midgut whose proper development depends on BMP signaling.^41,63^ We next asked whether the acid zone impairment observed in *Ostγ*-mutant larvae is linked to defects in embryonic Dpp signaling. To this end, *Drosophila* embryos were stained with an antibody against human pSMAD3, which recognizes *Drosophila* pMad,^64^ the effector of Dpp signaling in the signal-receiving cells.^65^ As shown in Figure 3D, heterozygous embryos (controls) showed pMad staining in areas corresponding to PS3 and PS7. However, *Ostγ ^Δ5/Δ5^* embryos showed a dramatic decrease in the level of pMad in PS3 and PS7. Notably, pMad staining in other regions of the embryos, including the ectodermal and head regions, was not affected by the loss of *Ostγ*. We further marked the precursors of the midgut acid zone cells in PS7 with anti-Labial antibody and the VM cells with anti-Fas3 (Fasciclin 3) antibody. As shown in Figure 3E, Labial expression was detected in the PS7 region of the control larvae at stage 14. However, in *Ostγ* mutant embryos, no Labial signal was detectable in the PS7 region. Together, these data indicate that loss of *Ostγ* leads to impaired BMP signaling in the *Drosophila* midgut.

To examine the impact of loss of *Ostγ* on the Dpp protein, we performed western blot on larval midguts with an anti-Dpp antibody, which recognizes the prodomain of Dpp.^66^ As shown in Figure 3F, Dpp is accumulated as a high molecular weight band in *Ostγ* mutant larvae (*Ostγ ^Δ5/Δ5^* and *Ostγ ^Δ5/Df^*) compared to controls (*y w*, and *Ostγ ^Δ5/+^*), suggesting retention of glycosylated species in the larval midgut as shown above in MEFs.

The shift in the Dpp migration pattern *in vivo* and our observations in MEFs suggest that TUSC3/Ostγ functions in the BMP4/Dpp signal-sending cells to regulate the BMP ligand. To test this notion *in vivo*, we performed mesodermal RNAi-mediated knockdown (KD) and rescue experiments. At stage 14, *Mef2>Ostγ^RNAi^* embryos showed a strongly decreased Labial expression in PS7 (Figure 4A) and a full impairment of the acid zone in the larval midgut (Figure 4B) when compared to the control larvae. These phenotypes are accompanied by a partially penetrant shortening of gastric caeca (Figure 4B). These observations strongly suggest that *Ostγ* is required in Dpp-expressing VM cells to promote BMP signaling. Although ubiquitous RNAi-mediated knockdown of *Ostγ* recapitulated the full lethality phenotype of mutants, we found different degrees of lethality with two mesodermal drivers (*Mef2-GAL4* and *how^24B^-GAL4*) and a neuronal driver (*Elav-GAL4*), suggesting that *Ostγ* also plays key roles in other tissues and organs beyond the embryonic and larval midgut (Figure 4C).

**Figure 4.**
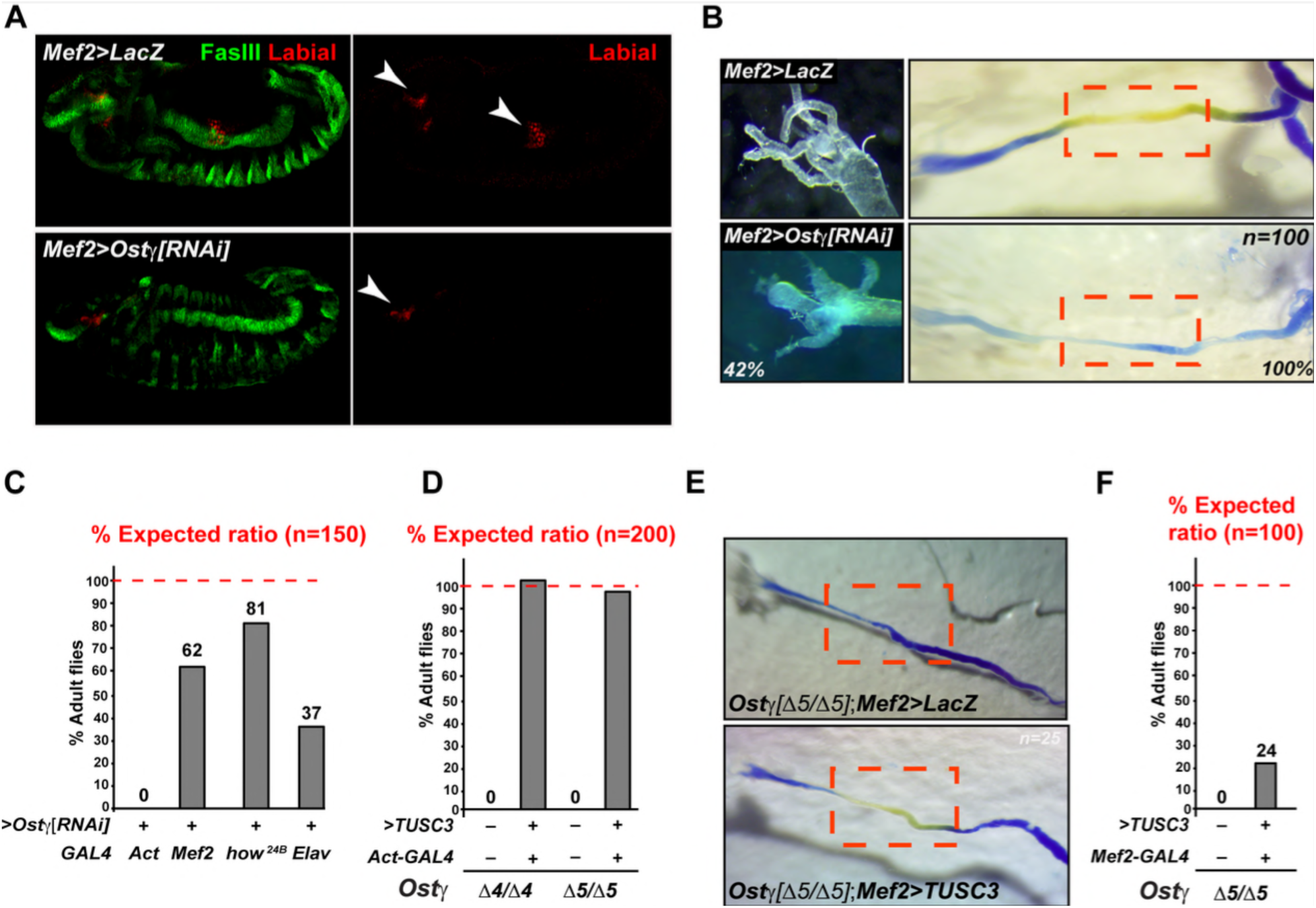
Mesodermal *Ostγ* is necessary for VM-specific Dpp signaling and is functionally rescued by human *TUSC3*. (A). Stage-14 embryos expressing *Ostγ^RNAi^* under *Mef2-GAL4 (Mef2>Ostγ^RNAi^)* or *GAL4* alone were stained for Labial (red) and Fas3 (green). Arrowheads mark PS3 and PS7. (B) Representative gut images of third-instar larvae expressing *Mef2>Ostγ^RNAi^ or Mef2>+* and fed with BPB. Dashed boxes mark the acid zone area. Scale bar, 200 µm. (C and D) Survival curves plotting percentage of adult eclosion from the indicated genotypes. Numbers indicate percent survival. (E) Representative midgut images of third-instar larvae with the indicated genotypes fed with BPB. (F) Survival curves plotting percentage of adult eclosion from the indicated genotypes.

To directly examine the degree of functional conservation between fly *Ostγ* and TUSC3, we generated transgenic flies capable of overexpressing human *TUSC3* using *ΦC31*-mediated transgenesis.^67,68^ When crossed with a ubiquitous driver (*Act-GAL4*), the *TUSC3* overexpressing line was able to fully rescue the lethality of both *Ostγ* alleles (Figure 4D). Moreover, its mesodermal overexpression restored the acid zone in *Ostγ ^Δ5/Δ5^* mutant larvae (Figure 4E). Notably, in accordance with the partial lethality of *Ostγ* mesodermal KD, *TUSC3* overexpression with *Mef2-GAL4* partially rescued the *Ostγ ^Δ5/Δ5^* lethality (Figure 4F). Together, these results demonstrate that human *TUSC3* can compensate for the loss of *Ostγ*, strongly suggesting that it promotes endogenous Dpp signaling in flies similar to Ostγ.

### TUSC3 functions as a rate-limiting gatekeeper of ER-folding and Dpp/BMP4 secretion

Our data strongly suggest that TUSC3 has a role in the early stage of Dpp/BMP4 glycosylation. To test whether the level of TUSC3 modulates the ER-folding state of Dpp/BMP4 molecules, *Tusc3* was overexpressed and compared with control and *Tusc3* knockdown MEFs. As shown in Figure 5A, the OS9 and BiP lectin-chaperones were found less intense in *Tusc3* knockdown MEFs, suggesting that the accumulation of G2M9-BMP4 is not sensed by the ERAD machinery. However, upon *Tusc3* overexpression, the OS9 and BiP lectin-chaperones also showed a decline in expression when compared to control MEFs. Surprisingly, although the activation of the OS9 and BiP lectin-chaperones displayed a similar trend in *Tusc3* overexpression and knockdown experiments, anti-HA immunoblotting showed that the G2M9-BMP4-HA band is strongly decreased by *Tusc3* overexpression, as well as the deglycosylated BMP4-HA band. Moreover, secretion of BMP4-Myc in the media was increased upon *Tusc3* overexpression, and this was accompanied by increased pSMAD1/5 level, which indicate enhanced BMP signaling. These observations suggest that TUSC3 serves as a rate-limiting checkpoint in early *N*-glycoprotein maturation: *Tusc3* knockdown leads to the accumulation of glucosylated BMP4 molecules which fail to engage ERQC, whereas its overexpression licenses BMP4’s G2M9 to G1M9 trimming and promotes its trafficking through the secretory pathway, thereby enhancing BMP signaling.

**Figure 5.**
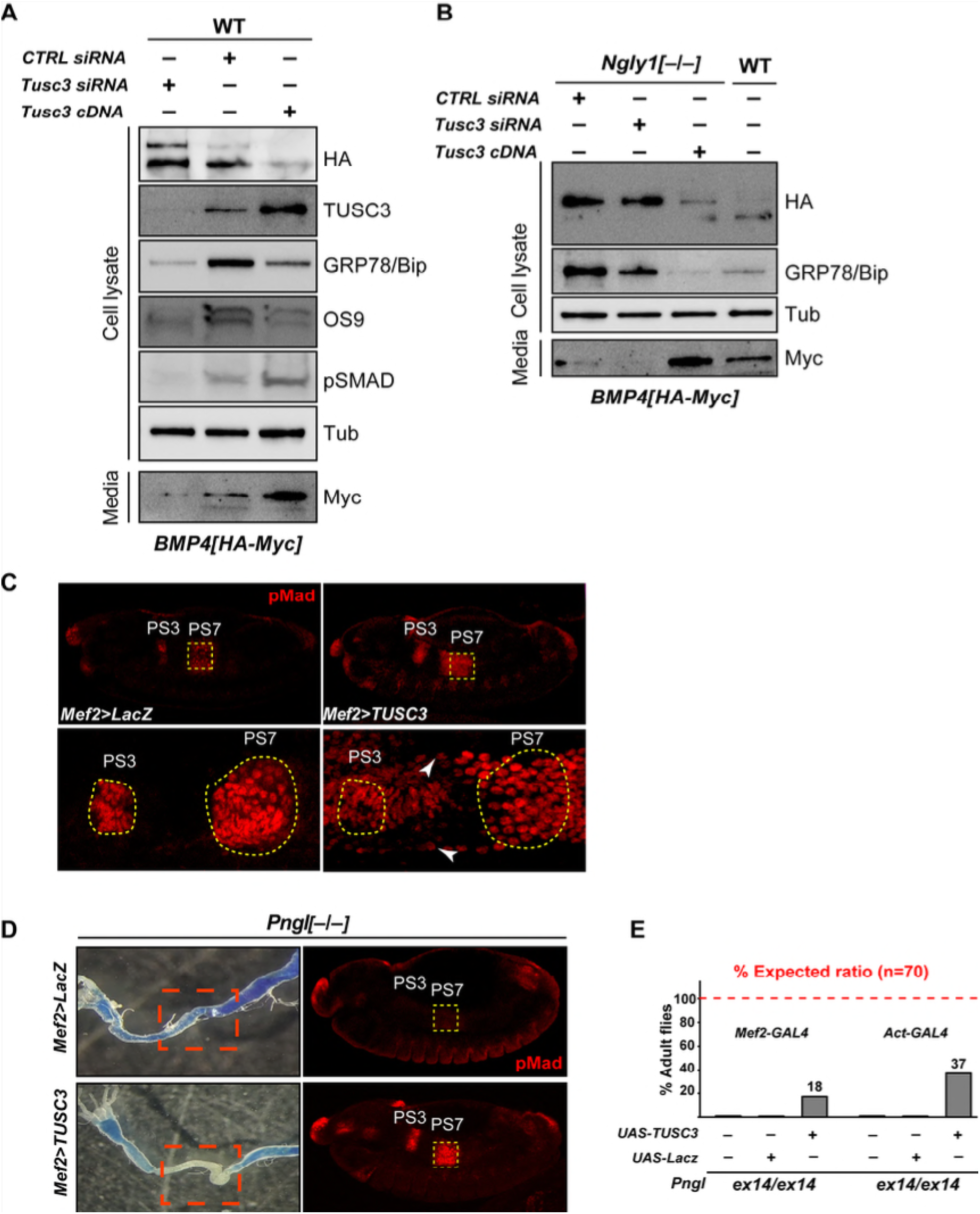
TUSC3 is a rate-limiting regulator of BMP4 folding, secretion, thereby affecting ERAD degradation. (A) Western blot of BMP4-HA and BMP4-Myc in cell lysates and media from control, *Tusc3*-knockdown, and *Tusc3*-overexpressing MEFs. BiP and OS9 included as ER stress markers. Tubulin serves as loading control. (B) Western blot analysis of BMP4-HA, BMP4-Myc, and BiP in wild-type and *Ngly1^−/−^* MEFs under control, *Tusc3-*knockdown, and *Tusc3-*overexpression conditions. Representative of two independent experiments. (C) Immunofluorescence staining of pMad in stage 14 embryos under mesodermal *TUSC3* overexpression. Dashed shapes mark PS3 and PS7; white arrowheads mark the expandion of the pMad⁺ domain beyond PS3/PS7. (D) Acid zone (BPB assay) and pMad staining in *Pngl^−/−^* embryos (Stage-14 embryos) with or without mesodermal *TUSC3* overexpression. Representative images of 20 midguts and 5 embryos. (E) Survival curves plotting percent adult eclosion of *Pngl^−/−^*progeny carrying *Mef2>TUSC3 or Act>TUSC3* compared to *Pngl* mutants.

The above observations prompted us to ask whether TUSC3 promotes proper folding of BMP4 molecules early during the quality control process, leading to an increase in the secretion-eligible molecules and a reciprocal reduction in misfolded BMP4 molecules that will need to undergo ERAD. As we previously reported, the cytosolic enzyme *N*-glycanase 1 (NGLY1) promotes the retrotranslocation of misfolded BMP4 molecules from the ER and their ERAD, thereby preventing the accumulation of misfolded BMP4 in the ER.^41^ Accordingly, we examined the effects of *Tusc3* knockdown and overexpression on BMP4 trafficking and BiP induction in *Ngly1^−/−^* MEFs. In agreement with our previous report, when compared to control MEFs, *Ngly1^−/−^* MEFs exhibited a strong accumulation of *N-*glycosylated BMP4 molecules, a loss of the de-*N*-glycosylated BMP4 molecules—as expected upon loss of the sole *N-*glycanase activity from the cells—and a reduction in the level of BMP4-Myc in media (Figure 5B). *Ngly1^−/−^* MEFs also showed an increase in BiP levels, as expected from accumulation of misfolded BMP4 in the ER. *Tusc3* knockdown did not affect the level of glycosylated BMP4 in the cell in *Ngly1^−/−^* MEFs but reduced BiP level and further reduced BMP4-Myc secretion in the media. In sheer contrast, overexpression of *Tusc3* in *Ngly1^−/−^* MEFs led to a strong reduction in the level of glycosylated BMP4-HA, a strong increase in the level of BMP4-Myc in the media, and a dramatic reduction of Bip levels, even compared to control cells overexpressing BMP4-HA-Myc (Figure 5B). Together, these data suggest that TUSC3 promotes proper folding of BMP4 molecules in a dosage-sensitive manner, thereby reducing the number of BMP4 molecules selected for ERAD and NGLY1-mediated deglycosylation and instead increasing the number of BMP4 molecules that traffic through the secretory pathway, ultimately leading to increased secretion of active (cleaved) BMP4.

Our conclusions from the above-mentioned TUSC3 overexpression experiments are based on data generated in a BMP4 overexpression context. Therefore, we sought to examine whether overexpression of *TUSC3* in the *Drosophila* larval VM can increase BMP signaling mediated by endogenous Dpp. Indeed, overexpression of *TUSC3* in the mesoderm led to an expansion of pMad^+^ domain beyond the PS3 and PS7 regions in the embryonic midgut compared (Figure 5C), suggesting a gain of Dpp signaling. To further examine whether the gain of signaling directly depends on increasing Dpp signaling, we overexpressed *TUSC3* in the VM of *Pngl^−/−^*flies. *Pngl,* which is the functional ortholog of *NGLY1* in *Drosophila*, promotes Dpp signaling in the embryonic midgut by enzymatically removing *N-*glycans from misfolded Dpp proteins in the ER, facilitating their ERAD and allowing the properly folded Dpp molecules to be secreted by the VM cells.^34,41^ As reported, loss of *Pngl* impaired the formation of acid zone in the larval midgut due to the loss of Dpp signaling, as evidenced by the lack of pMad^+^ cells in the PS7 region (Figure 5D, top panels; compare to control larvae in Figure 3C)). When *TUSC3* was overexpressed in Dpp sending cells of *Pngl* mutant midgut, the larval acid zone and pMad staining in the PS7 region in the embryonic midgut were fully restored (Figure 5D, bottom panels). Notably, *TUSC3* overexpression in the mesoderm of *Pngl* mutant larvae led to ∼ 18% rescue of *Pngl* lethality (Figure 5E), which is comparable to the 22% rescue of *Pngl^−/−^* lethality in animals in which three of the Dpp *N*-glycosylation sites are ablated.^41^ These results confirm our observations in MEFs and strongly indicate that TUSC3 promotes proper folding of Dpp molecules in the ER of the VM cells, leading to enhanced BMP signaling in neighboring endodermal cells and reduces the number of misfolded Dpp molecules in these cells, ultimately suppressing of the *Pngl* loss-of-function phenotype in this tissue. Importantly, the level of lethality rescue was doubled when *TUSC3* was ubiquitously expressed in *Pngl^−/−^ Drosophila*, indicating that TUSC3’s role in promoting protein folding upstream of ERAD entry and NGLY1-mediated deglycosylation is not limited to BMP signaling in embryonic mesodermal tissue.

## DISCUSSION

Our study identifies TUSC3 as a conserved, dosage-sensitive regulator of a previously unrecognized early ER triage checkpoint that governs the fate of BMP4/Dpp. Stepwise glucose trimming is a well-established aspect of *N*-glycosylation, wherein the first glucose is believed to be constitutively removed, and the third glucose enables ERQC via calnexin binding.^3,16^ The functional relevance of the second glucose, however, had remained undefined. Our data indicate that TUSC3 engages nascent BMP4/Dpp and promotes GCS2-mediated removal of the second glucose from high-mannose glycans on these proteins as a rate-limiting step for entry into the calnexin/calreticulin-dependent ERQC. This conclusion is supported by the following key observations (Figure 6): (1) Upon loss of *TUSC3*, G2M9-decorated BMP4/Dpp molecules fail to transition to G1M9, aggregate in the ER, and evade both UPR and ERAD, resulting in impaired secretion and diminished pSMAD/Mad signaling; (2) Malectin binding analyses confirm that the BMP4 molecules accumulated in *Tusc3*-deficient cells bear high-mannose G2M9 glycans; (3) ER stress markers such as BiP and pIRE1α are not induced in TUSC3-deficient cells, indicating escape from canonical UPR surveillance; (4) Overexpression of TUSC3 accelerates G2M9 to G1M9 glycan trimming, shifting BMP4 cargo into productive folding and secretion and enhancing BMP signaling. Importantly, loss- and gain-of-function studies in *Drosophila* corroborate the role of *TUSC3* in regulating ER triage of the endogenously expressed Dpp (the fly homolog of BMP4).

**Figure 6.**
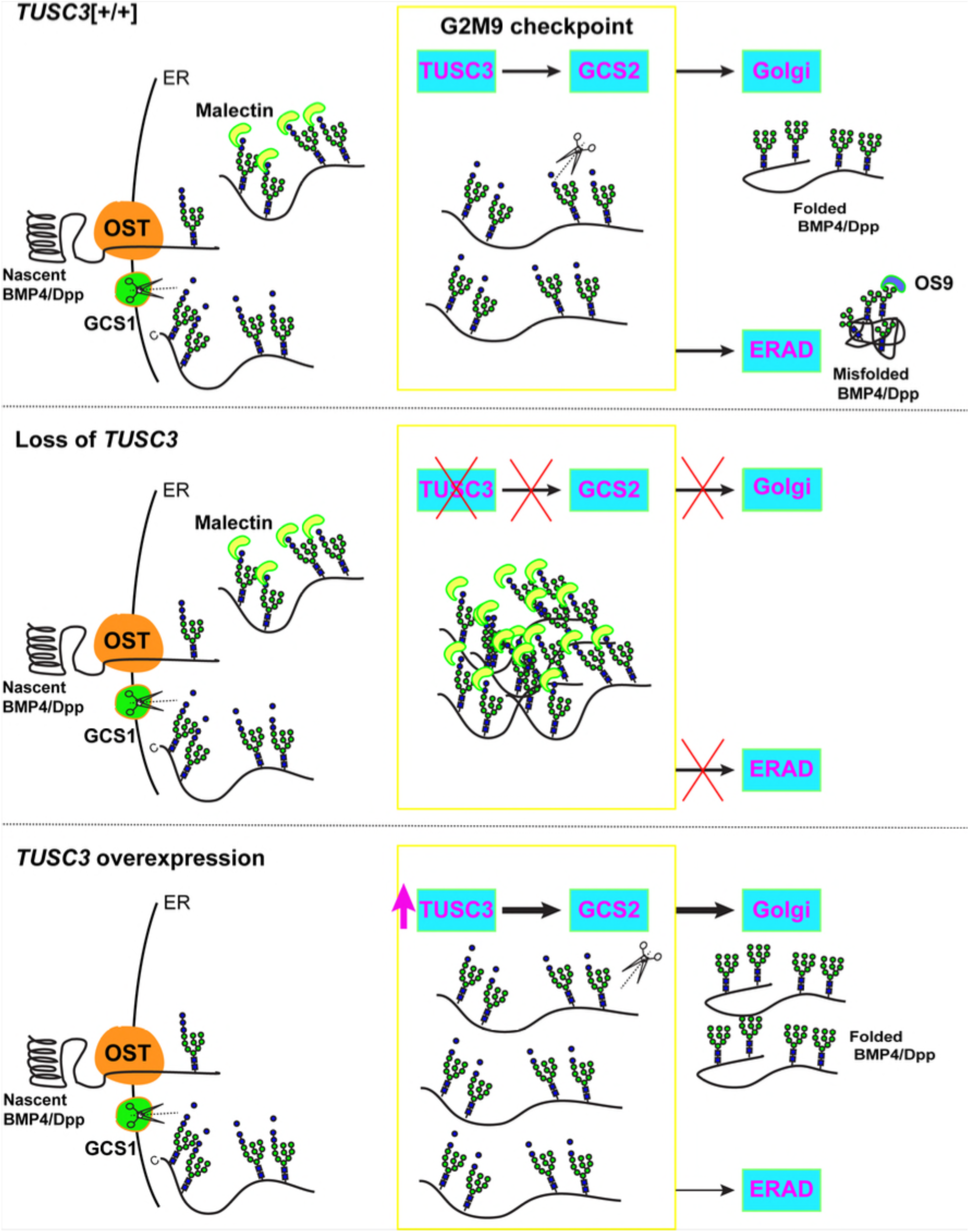
Model of TUSC3-mediated G2M9 checkpoint in ER quality control and BMP4/Dpp secretion. Schematic of the function of TUSC3/Ostγ as a rate-limiting gatekeeper for nascent *N-*glycosylated BMP4/Dpp molecules in the ER. TUSC3 (or fly *Ostγ*) facilitates G2 to G1 trimming of G2M9-BMP4 by ER α-glucosidase II (GCS2). Once trimmed to G1M9, the glycoprotein can proceed through productive folding and assembly into secretion-competent BMP4 homodimers. In the absence of *TUSC3/Ostγ*, G2M9-decorated BMP4/Dpp molecules accumulate in the ER as insoluble aggregates that bind malectin but fail to recruit calnexin/calreticulin or recruit OS9 and BiP. These aggregates evade pIRE1α/BiP induction and ERAD engagement, leading to impaired secretion. When *TUSC3/Ostγ* is overexpressed, enhanced G2 to G1 trimming reduces the pool of di-glucosylated intermediates and additional BMP4/Dpp molecules are routed towards productive folding and away from ERAD.

Moreover, a malectin-positive BMP4 population is readily detected under basal conditions, indicating that G2M9-BMP4 is not merely an abnormal byproduct of *Tusc3* knockdown and GCS2 inhibition but is a physiological glycoform appearing during BMP4 maturation. Together, these findings underscore the physiological relevance of the TUSC3-regulated G2M9 checkpoint in this process. More broadly, OST subunit composition and dosage likely calibrate pathway output not only via site-specific occupancy but also by constraining the throughput of the early G2 to G1 licensing step, thereby limiting entry into ERQC.

While TUSC3 is not classically considered an ERAD component, emerging data suggest that it indirectly influences ERAD. For example, it has been proposed that TUSC3 regulates ERAD susceptibility through disulfide bond modulation or by altering glycoprotein folding kinetics.^9,69,70^ Our data indicate that TUSC3 acts upstream of retrotranslocation to divert BMP4 and potentially other substrates away from ERAD and toward productive folding. The most compelling evidence for this notion is the ability of TUSC3 overexpression to rescue BMP4 secretion and reduce ER stress in *Ngly1^−/−^* MEFs—where ERAD is impaired due to the lack of BMP4 retrotranslocation ^41^— and to restore BMP signaling and partial rescue the lethality in *Ngly1*-null flies (*Pngl^−/−^*). How does TUSC3 favor proper folding and secretion of BMP4? Our data support two scenarios which are not mutually exclusive: (1) directly facilitating GCS2-mediated trimming of the second glucose from G2M9 glycoform to promote and potentially accelerate the ERQC entry of the resulting G1M9-BMP4 molecules, thereby improving their folding; and/or (2) promoting substrate folding, potentially by preventing the aggregation G2M9-BMP4 molecules and facilitating their interaction with ER chaperones to render the G2–G1 bond in G2M9 accessible for trimming by GCS2. Regardless of the precise mechanism, these observations indicate that the pool of BMP4 molecules that normally undergoes ERAD is not irreversibly misfolded and can be shifted to a signaling-competent pool through a TUSC3-GCS2-mediated mechanism. In this scenario, the OST–GCS2– ERQC module has a limited capacity, which sets an upper limit on how many ligands can fold correctly and be secreted at a given time.

TUSC3 and MAGT1 possess a thioredoxin-like domain with a conserved CXXC motif, positioned adjacent to a peptide-binding groove that can accommodate substrates. Structural studies have proposed that this domain may facilitate STT3B-mediated glycosylation by forming transient disulfide bonds with cysteine-rich polypeptides.^21,22,46^ Notably, the mature domain of BMP4 and its fly homolog Dpp contains a highly conserved *N-*glycosylation sequon near key cysteine residues critical for intramolecular disulfide bonding and dimerization,^40,43^ raising the possibility that thioredoxin-mediated interactions could promote the maturation of BMP4/Dpp. However, work in MAGT1-deficient models of XMEN disease showed that mutating the CXXC motif did not impair MAGT1’s ability to rescue glycosylation of NKG2D, suggesting that disulfide crosslinking may not be essential for glycosylation of at least some substrates. Whether TUSC3’s thioredoxin domain contributes directly to BMP4/Dpp folding and their glucose trimming by GCS2 remains to be determined.

Our findings have implications for understanding the pathophysiology of TUSC3-CDG, a human disease caused by recessive *TUSC3* mutations and characterized by intellectual disability.^71–73^ The molecular basis for the neurological manifestations of these patients is not known, although defects in Mg^2+^ homeostasis have been suggested as a potential contributor.^30,31^ Our work suggests that impaired BMP signaling, due to defective G2M9 triage, could contribute to neurodevelopmental defects in patients with TUSC3-CDG. In addition, the observed restoration of BMP signaling in NGLY1-deficient MEF and flies by TUSC3 overexpression might offer an unexpected therapeutic strategy for diseases like NGLY1-CDG, which impair the ERAD of glycoproteins like BMP4. Beyond impaired neurodevelopment, *TUSC3* downregulation has been associated with tumorigenesis in multiple cancers.^23,26^ In prostate cancer models, *TUSC3* knockdown reduces PERK–CHOP and IRE1α–XBP1 activation while enhancing ATF6α signaling, leading to attenuated sensitivity to tunicamycin-induced stress.^26,30^ Similarly, *TUSC3* deficiency in A549 non–small cell lung cancer cells alters the UPR response with a decrease in IRE1 and PERK signaling while enhancing ERAD via the ATF6–HRD1 axis, thereby promoting metastatic potential.^26^ Differences in ERAD outcomes across TUSC3-deficent models may reflect variations in the level and glycosylation pattern of its glycoprotein clients and/or chaperone expression levels. Our data reveal that loss of *TUSC3* leads to silent accumulation of G2M9-decorated BMP4 aggregates that evade canonical UPR sensors such as BiP and pIRE1α. It remains to be seen whether this failure to engage ER stress sensors can help explain the reported tumor-suppressive role of TUSC3, particularly in contexts where misfolded glycoproteins escape degradation and accumulate without activating ER stress responses.

### Limitations of the study

While our findings establish TUSC3 as a conserved regulator of a G2M9-dependent ER triage checkpoint, several mechanistic and physiological aspects remain unresolved. First, the molecular mechanism by which TUSC3 facilitates GCS2-mediated G2 to G1 trimming is not fully defined. It remains unclear whether TUSC3 physically interacts with GCS2, functions by spatially positioning G2M9-decorated substrates, and/or promotes the folding of the nascent protein in a manner that allows GCS2 to recognize the G1–G2 bond. Second, we previously reported that during *Drosophila* development, Ngly1/Pngl functions in the mesoderm to regulate BMP signaling in a tissue-specific manner. Therefore, the observation that ubiquitous expression of human TUSC3 is twice as effective as its mesodermal expression in rescuing the lethality of *Pngl^−/−^*flies suggests that targets other than BMP4 are also subject to TUSC3’s regulatory role upstream of ERAD described here. Finally, although TUSC3 is known to participate in the STT3B-containing OST complex,^74^ our study does not allow us to determine whether TUSC3 specifically promotes G2M9 glucose trimming of *N-*glycans that are added post-translationally, or whether its activity also extends to co-translationally transferred glycans. Furthermore, we cannot exclude that TUSC3 preferentially regulates glycans located near the C-terminus of substrates. Recent works showed that the position of *N-*glycans acts as part of a “glyco-code” that influences the sequence of chaperone engagement, with *N-*terminal glycans favoring early lectin binding, whereas *C-*terminal glycans often rely on BiP assistance and subsequent re-glucosylation by UGGT to enable late lectin engagement^5^. Whether TUSC3 preferentially regulates such *C-*terminal glycans, as suggested by this model, remains to be shown.

## RESOURCE AVAILABILITY

### Lead contact

Requests for further information and resources should be directed to and will be fulfilled by the lead contact, Antonio Galeone.

### Materials availability

Newly generated plasmids and fly lines will be provided upon request under a Materials Transfer Agreement (MTA).

### Data and code availability

No new code was generated. Raw immunoblots and confocal images are available upon request.

## ACKNOWLEDGMENTS

We thank Angelo Santino and Pietro Roversi for discussions; Elisabetta Cuna and Andrea Lia for excellent technical assistance; The Bloomington *Drosophila* Stock Center, Jan Christian, and Tadashi Suzuki for reagents. This work was supported by grants from H2020-MSCA individual fellowship (844147) and Buzzati-Traverso fellowship (to AG and TV), young researchers MSCA grant, funded by the Italian Ministry of University and Research (MUR) through NextGenerationEU program (to AG), MUR PNRR “National Center for Gene Therapy and Drugs based on RNA Technology”, “Tecnopolo per la medicina di precisione” (TecnoMed Puglia) - Regione Puglia (to GG), NIH (R35GM130317) and Grace Science Foundation (to HJN). TV also acknowledges the support of the Italian ministry of research (PRIN grant 2020CLZ5XW). Imaging was performed at Unitech NOLIMITS microscopy facility core of University of Milan and Imaging facility at CNR Nanotec.

## AUTHOR CONTRIBUTIONS

Conceptualization, A.G., H.J.N and T.V.; methodology, A.G., H.J.N and T.V.; validation, A.G., E.S., F.L, S.Y.H., B.M., R.R.; formal analysis, A.G., E.S., R.R.., H.J.N and T.V; investigation, A.G., E.S., F.L, S.Y.H., B.M.; resources, A.G., H.J.N and T.V.; writing – original draft,. A.G., E.S., H.J.N and T.V..; writing – review & editing, all authors; project administration, A.G. and T.V.; funding acquisition, A.G., G.G., H.J.N and T.V

## STAR★METHODS

### KEY RESOURCES TABLE

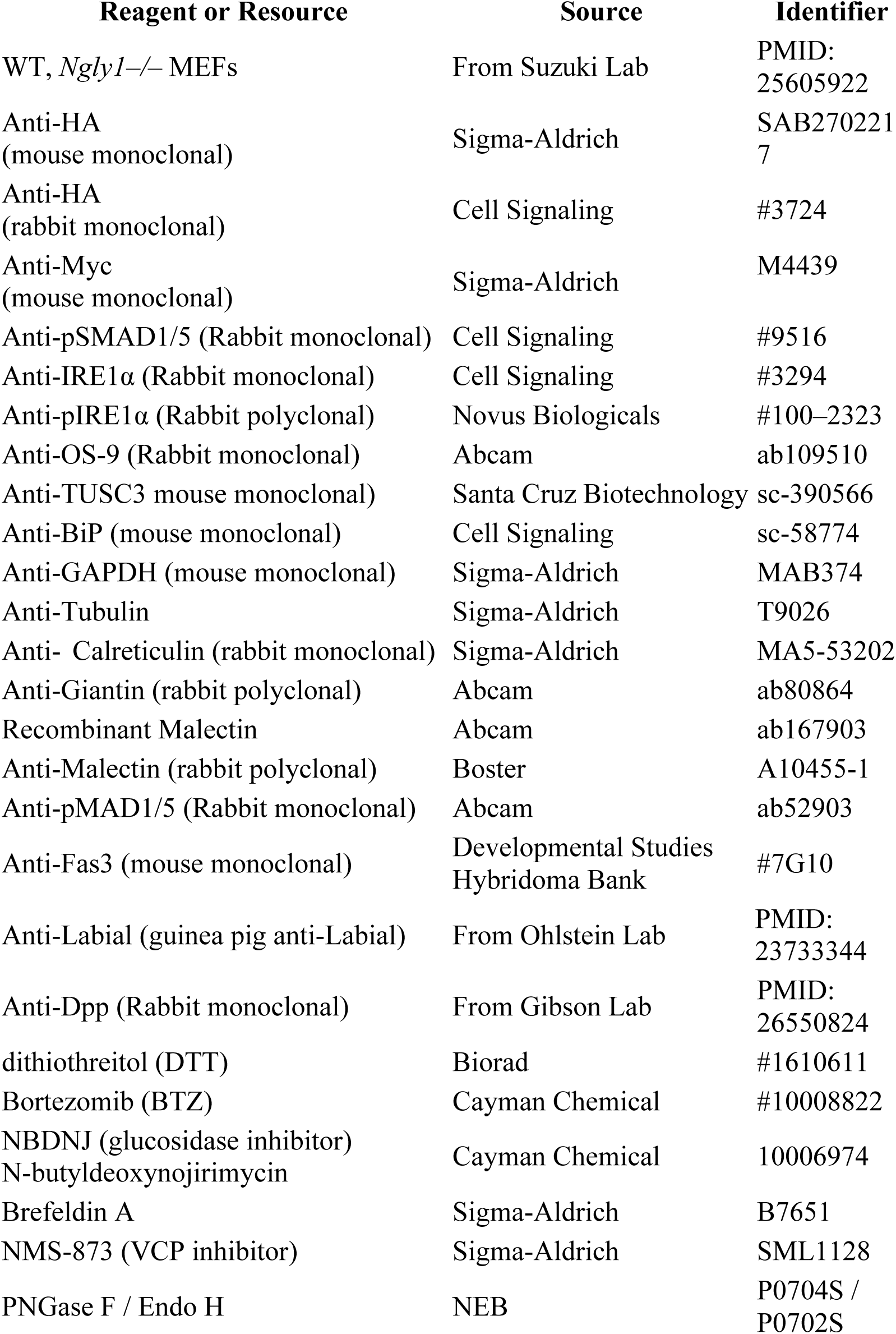
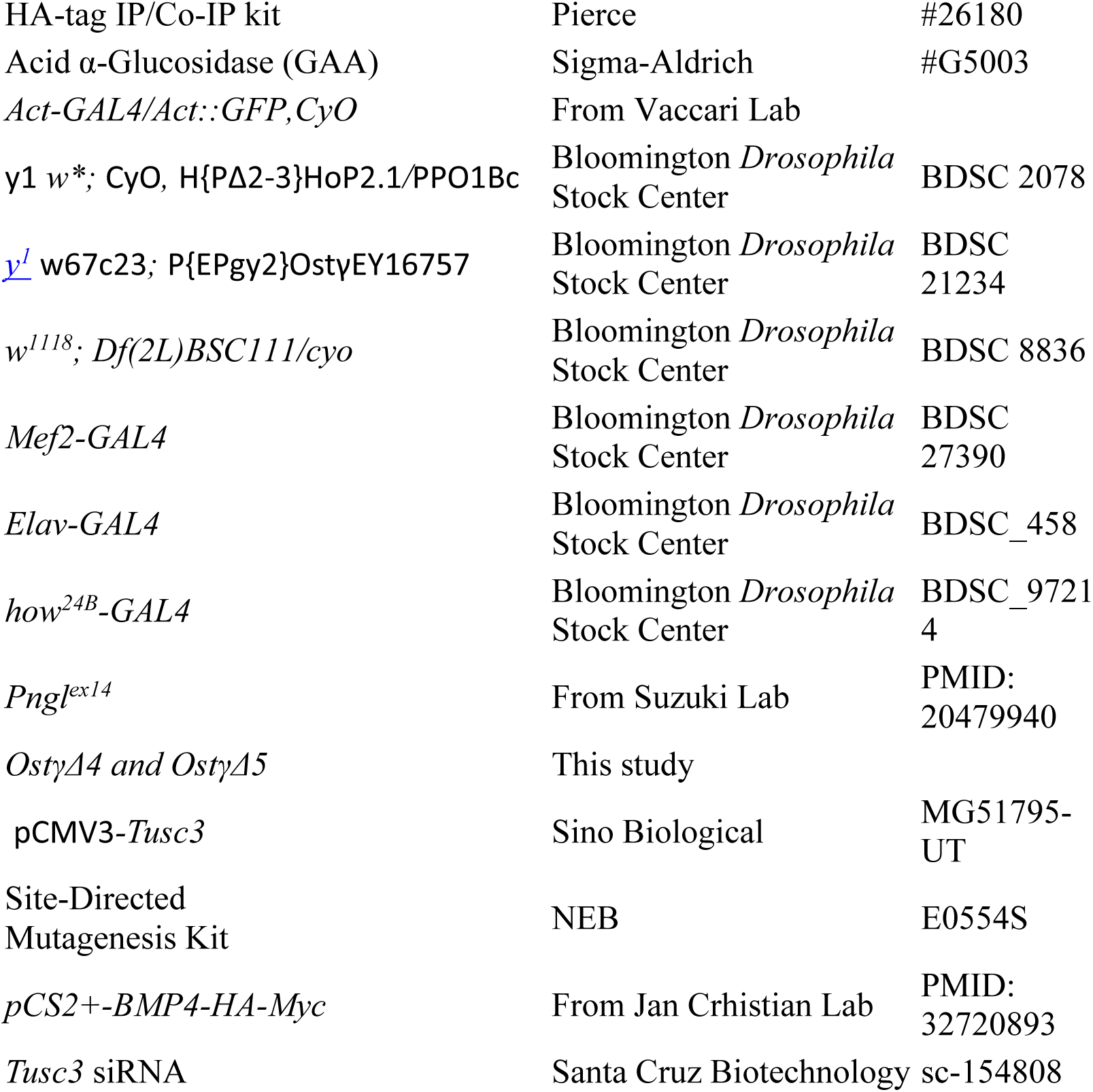

#### Mouse Embryonic Fibroblasts (MEFs)

Mouse embryonic fibroblasts (MEFs; WT and *Ngly1–/–*) were maintained in Dulbecco’s Modified Eagle Medium (DMEM; Sigma-Aldrich) supplemented with 10% fetal bovine serum (FBS; Gibco) and 100 U/mL penicillin–100 µg/mL streptomycin (Sigma-Aldrich) at 37 °C in 5% CO₂. For BMP4-HA-Myc expression, MEFs were transfected with *pCS2+-BMP4-HA-Myc* using Lipofectamine 3000 (Thermo Fisher Scientific) according to the manufacturer’s protocol. Small interfering RNA (siRNA) targeting mouse *Tusc3* (*siTusc3*) or non-targeting control (siCtrl) was transfected at 50 nM final concentration with Lipofectamine 3000. For overexpression of *Tusc3*, MEFs were transfected with a plasmid encoding full-length murine *Tusc3* cDNA (Tusc3-CDS) or empty vector using Lipofectamine 3000. In all experiments, transfection efficiency was monitored by immunoblotting using TUSC3.

##### Drosophila melanogaster

Flies were reared at 25 °C on standard cornmeal agar. Embryos were staged, fixed, and stained at developmental stage 14.

### METHOD DETAILS

#### BMP4 Glycoforms: protein extraction, immunoprecipitation, and western Blotting

MEFs were washed twice with ice-cold PBS and lysed in RIPA buffer (50 mM Tris-HCl, pH 7.6; 150 mM NaCl; 1% NP-40; 0.1% SDS; 0.5% sodium deoxycholate) supplemented with protease inhibitor cocktail (Roche) for 30 min on ice. Lysates were clarified by centrifugation at 14,000 × g for 10 min at 4 °C, and supernatants were collected for total protein quantification (BCA assay; Pierce). Conditioned media were collected, cleared by centrifugation at 1,000 × g for 5 min, and precipitated with 10% (w/v) trichloroacetic acid (TCA)/acetone overnight at –20 °C. Protein pellets were washed once with acetone, air-dried, and resuspended in 1× Laemmli buffer.

For immunoprecipitation (IP), 500 µg total protein from MEF lysates was incubated with 30 µL anti-HA magnetic beads (Pierce HA-Tag IP Kit) overnight at 4 °C with rotation. Beads were washed three times with NP-40 lysis buffer (50 mM Tris-HCl, pH 7.5; 150 mM NaCl; 1% NP-40; 10% glycerol; protease inhibitors), and IP products were eluted with 1× Laemmli buffer by boiling at 95 °C for 5 min. Equal amounts of protein (20–30 µg) from cell lysates or TCA-precipitated media were resolved on 7.5% SDS–PAGE gels (Bio-Rad) and transferred to nitrocellulose membranes (0.45 µm; Bio-Rad) using the Trans-Blot Turbo system (Bio-Rad). Membranes were blocked in 5% (w/v) nonfat dry milk in TBST (20 mM Tris-HCl, pH 7.6; 150 mM NaCl; 0.1% Tween-20) for 1 h at room temperature and incubated with primary antibodies overnight at 4 °C. After primary incubation, membranes were washed three times in TBST and incubated with species-appropriate HRP-conjugated secondary antibodies for 1 h at room temperature. Blots were developed with ECL substrate (Pierce) and imaged on a ChemiDoc MP System (Bio-Rad).

For deglycosylation assays, 20 µg of MEF lysate was denatured in 1× denaturing buffer (0.5% SDS, 40 mM DTT; New England Biolabs) for 10 min at 100 °C. Samples were cooled to room temperature, and GlycoBuffer 3 (50 mM sodium phosphate, pH 7.5) was added to 1× final concentration. PNGase F or Endo H was added, and reactions were incubated at 37 °C for 1 h. The reaction was terminated by adding 4× Laemmli buffer and heating at 95 °C for 5 min. Digested samples were analyzed by SDS–PAGE and immunoblotting as described above. Malectin far-western blotting was used to probe G2M9 glycoforms.

For the aggregation assay, MEFs were lysed in NP-40 digestion buffer (50 mM Tris–HCl, pH 7.5; 150 mM NaCl; 1 mM EDTA; 1% Triton X-100; protease inhibitors) on ice for 30 min. Lysates were centrifuged at 14,000 × g for 10 min at 4 °C. The supernatant (soluble fraction) was separated from the pellet (insoluble fraction). Pellets were washed once in NP-40 buffer, resuspended in 1× Laemmli buffer, and sonicated briefly and then treated with urea (4 and 8M). Equal proportions of soluble and insoluble fractions were analyzed by SDS–PAGE and immunoblotting for anti–HA.

#### Inhibitor Treatments

To induce ER stress, MEFs were treated with 2 mM dithiothreitol for 30 min at 37 °C. Proteasome inhibition was achieved by incubating cells with 10 nM bortezomib (BTZ) or vehicle (0.1% DMSO) for 6 h. For VCP inhibition, cells were treated with 5 µM NMS-873 for 6 h. Brefeldin A was used at 1 µg/mL for 6 h to block ER–Golgi trafficking. To inhibit ER α-glucosidases I and II, MEFs were incubated with 2 mM N-butyldeoxynojirimycin for 8 h. For cold-temperature ER enrichment, cells were shifted to 20 °C for 4 h prior to harvest.

##### *Drosophila* Embryo Staining and Viability

Embryos were collected on grape juice agar plates supplemented with yeast paste, dechorionated in 50% bleach for 3 min, rinsed in PBS, and fixed in 4% paraformaldehyde/heptane for 20 min at room temperature. Fixed embryos were devitellinized by shaking in methanol and stored in methanol at –20 °C. For immunostaining, embryos were rehydrated through a methanol/PBST (0.1% Tween-20 in PBS) series and blocked in PBST + 5% normal goat serum (NGS) for 1 hour.

For survival (eclosion) tests, the expected ratio of offspring was calculated based on Mendelian inheritance for each genotypic class and the observed/expected ratio is reported as a percentage.

For midgut clearance assay, third-instar larvae were dissected in cold PBS and transferred to food containing 0.025% (w/v) bromophenol blue (BPB; Sigma-Aldrich) for 2 h at 25 °C. Midguts were removed, briefly rinsed in PBS, and imaged using an Olympus SZX16 stereomicroscope. Presence of an acidified region (yellow BPB staining) was scored in at least 30 larvae per genotype.

#### Generation of *Ostγ* Imprecise Excision Alleles

A P-element insertion *(P{EPgy2}Ostγ^EY16757)* located in the 5′ UTR of the *Ostγ* locus (chromosome 2L:8,308,166) was mobilized by *Δ2-3* transposase to produce imprecise excisions. Virgin females carrying *P{EPgy2}Ostγ^EY16757* were crossed to males of the *Δ2-3, Sb/TM6, Ubx* stock (Bloomington Drosophila Stock Center [BDSC] #2078) at 25 °C. Progeny bearing both the *P-element* and *Δ2-3* source were selected by eye color and Sb marker, then intercrossed in groups of 10–15 to enrich for transposition events. Adult F₁ males exhibiting loss of the white^+ marker (indicative of P-element excision) were crossed individually to *w^1118; CyO/Sco* virgin females. F₂ siblings were screened for new deletions by PCR using genomic DNA extracted from single adults. Primers flanking the original insertion site (5′-GTCAGCGCTCTGTTGTTGAA-3′ and 5′-ACAGGAGAAACGTCCACGAG-3′) were used to amplify a ∼1.2 kb fragment in wild-type, with shorter products indicating imprecise excisions. PCR products from candidate deletion lines were subcloned (pCR™-Blunt II-TOPO®, Thermo Fisher) and Sanger-sequenced to map deletion breakpoints. Two alleles, *OstγΔ4 and OstγΔ5*, were selected based on non-overlapping internal deletions that remove coding exon sequences and are predicted to be null. Each allele was balanced over CyO; lethal phenotypes were confirmed by failure of homozygotes to eclose at 25 °C. For all downstream experiments, homozygous mutants were distinguished from balanced siblings by the absence of the *CyO* curly-wing marker.

#### Immunofluorescence and Confocal Microscopy (MEFs)

Glass-bottom dishes (MatTek) or coverslips were coated with poly-D-lysine (Sigma-Aldrich) and seeded with MEFs (50,000 cells/dish). After 24 h, cells were fixed in 4% paraformaldehyde (Electron Microscopy Sciences) in PBS for 15 min at room temperature, permeabilized in 0.2% saponin (Sigma-Aldrich) in PBS for 10 min and blocked in 5% normal goat serum (Jackson ImmunoResearch) for 1 h. Primary antibodies were diluted in blocking buffer and incubated overnight at 4 °C. After three washes in PBS, Alexa Fluor 488– or 594–conjugated secondaries were applied for 1 h at room temperature. Nuclei were counterstained with DAPI Samples were mounted in ProLong Gold Antifade and imaged on a Zeiss LSM 980 confocal microscope (63×/1.4 NA oil objective).

### QUANTIFICATION AND STATISTICAL ANALYSIS

Data represent mean ± SEM or mean ± SD from ≥3 biological replicates. Band intensities were normalized to loading controls. Densitometric quantification was performed with Fiji (ImageJ). Comparisons were analyzed using unpaired Student’s t-test or one-way ANOVA with Tukey post hoc correction (GraphPad Prism). Statistical significance: *p* < 0.05. Colocalization analysis (Pearson’s correlation coefficient) was performed in Fiji (ImageJ from three independent experiments.

## REFERENCES

1. Varki, A. (2017). Biological roles of glycans. Glycobiology 27, 3–49. 10.1093/glycob/cww086.

2. Schwarz, F., and Aebi, M. (2011). Mechanisms and principles of N-linked protein glycosylation. Curr Opin Struct Biol 21, 576–582. 10.1016/J.SBI.2011.08.005.

3. Helenius, A., and Aebi, M. (2004). Roles of N-linked glycans in the endoplasmic reticulum. Preprint, 10.1146/annurev.biochem.73.011303.073752 https://doi.org/10.1146/annurev.biochem.73.011303.073752.

4. Kanapin, A., Batalov, S., Davis, M.J., Gough, J., Grimmond, S., Kawaji, H., Magrane, M., Matsuda, H., Schönbach, C., Teasdale, R.D., et al. (2003). Mouse Proteome Analysis. Genome Res 13, 1335–1344. 10.1101/gr.978703.

5. Guay, K.P., Ke, H., Canniff, N.P., George, G.T., Eyles, S.J., Mariappan, M., Contessa, J.N., Gershenson, A., Gierasch, L.M., and Hebert, D.N. (2023). ER chaperones use a protein folding and quality control glyco-code. Mol Cell 83, 4524–4537.e5. 10.1016/J.MOLCEL.2023.11.006.

6. Adams, B.M., Oster, M.E., and Hebert, D.N. (2019). Protein Quality Control in the Endoplasmic Reticulum. Protein J 38, 317–329. 10.1007/s10930-019-09831-w.

7. Guay, K.P., Chou, W.-C., Canniff, N.P., Paul, K.B., and Hebert, D.N. (2025). N-glycan-dependent protein maturation and quality control in the ER. Nat Rev Mol Cell Biol. 10.1038/s41580-025-00855-y.

8. Kelleher, D.J., and Gilmore, R. (2006). An evolving view of the eukaryotic oligosaccharyltransferase. Glycobiology 16, 47R–62R. 10.1093/glycob/cwj066.

9. Hebert, D.N., Garman, S.C., and Molinari, M. (2005). The glycan code of the endoplasmic reticulum: asparagine-linked carbohydrates as protein maturation and quality-control tags. Trends Cell Biol 15, 364–370. 10.1016/j.tcb.2005.05.007.

10. Ellgaard, L., and Helenius, A. (2003). Quality control in the endoplasmic reticulum. Nat Rev Mol Cell Biol 4, 181–191. 10.1038/nrm1052.

11. Barker, M.K., and Rose, D.R. (2013). Specificity of Processing &#x3b1;-Glucosidase I Is Guided by the Substrate Conformation: CRYSTALLOGRAPHIC AND IN SILICO STUDIES *. Journal of Biological Chemistry 288, 13563–13574. 10.1074/jbc.M113.460436.

12. Moremen, K.W., and Molinari, M. (2006). N-linked glycan recognition and processing: the molecular basis of endoplasmic reticulum quality control. Curr Opin Struct Biol 16, 592–599. 10.1016/j.sbi.2006.08.005.

13. Kelleher, D.J., Karaoglu, D., Mandon, E.C., and Gilmore, R. (2003). Oligosaccharyltransferase Isoforms that Contain Different Catalytic STT3 Subunits Have Distinct Enzymatic Properties. Mol Cell 12, 101–111. 10.1016/S1097-2765(03)00243-0.

14. Burda, P., and Aebi, M. (1999). The dolichol pathway of N-linked glycosylation. Biochimica et Biophysica Acta (BBA) - General Subjects 1426, 239–257. 10.1016/S0304-4165(98)00127-5.

15. Mohorko, E., Glockshuber, R., and Aebi, M. (2011). Oligosaccharyltransferase: The central enzyme of N-linked protein glycosylation. Preprint, 10.1007/s10545-011-9337-1 https://doi.org/10.1007/s10545-011-9337-1.

16. Aebi, M. (2013). N-linked protein glycosylation in the ER. Biochimica et Biophysica Acta (BBA) - Molecular Cell Research 1833, 2430–2437. 10.1016/j.bbamcr.2013.04.001.

17. Braunger, K., Pfeffer, S., Shrimal, S., Gilmore, R., Berninghausen, O., Mandon, E.C., Becker, T., Förster, F., and Beckmann, R. (2018). Structural basis for coupling protein transport and N-glycosylation at the mammalian endoplasmic reticulum. Science (1979) 360, 215–219. 10.1126/science.aar7899.

18. Mohorko, E., Owen, R.L., Malojčić, G., Brozzo, M.S., Aebi, M., and Glockshuber, R. (2014). Structural Basis of Substrate Specificity of Human Oligosaccharyl Transferase Subunit N33/Tusc3 and Its Role in Regulating Protein <em>N</em>-Glycosylation. Structure 22, 590–601. 10.1016/j.str.2014.02.013.

19. Li, F., Chaigne-delalande, B., Kanellopoulou, C., Jeremiah, C., and Lenardo, M.J. (2012). Signaling role for Mg2+ revealed by immunodeficiency due to loss of MagT1. Nature 475.

20. Matsuda-Lennikov, M., Biancalana, M., Zou, J., Ravell, J.C., Zheng, L., Kanellopoulou, C., Jiang, P., Notarangelo, G., Jing, H., Masutani, E., et al. (2019). Magnesium transporter 1 (MAGT1) deficiency causes selective defects in N-linked glycosylation and expression of immune-response genes. Journal of Biological Chemistry 294. 10.1074/jbc.RA119.008903.

21. Cherepanova, N.A., Shrimal, S., and Gilmore, R. (2014). Oxidoreductase activity is necessary for N-glycosylation of cysteine-proximal acceptor sites in glycoproteins. Journal of Cell Biology 206, 525–539. 10.1083/jcb.201404083.

22. Shrimal, S., Cherepanova, N.A., and Gilmore, R. (2015). Cotranslational and posttranslocational N-glycosylation of proteins in the endoplasmic reticulum. Semin Cell Dev Biol 41, 71–78. 10.1016/j.semcdb.2014.11.005.

23. Horak, P., Tomasich, E., Vaňhara, P., Kratochvílová, K., Anees, M., Marhold, M., Lemberger, C.E., Gerschpacher, M., Horvat, R., Sibilia, M., et al. (2014). TUSC3 Loss Alters the ER Stress Response and Accelerates Prostate Cancer Growth in vivo. Sci Rep 4, 3739. 10.1038/srep03739.

24. Feng, S., Zhai, J., Lu, D., Lin, J., Dong, X., Liu, X., Wu, H., Roden, A.C., Brandi, G., Tavolari, S., et al. (2018). TUSC3 accelerates cancer growth and induces epithelial-mesenchymal transition by upregulating claudin-1 in non-small-cell lung cancer cells. Exp Cell Res 373, 44–56. 10.1016/j.yexcr.2018.08.012.

25. Ren, Y., Deng, R., Cai, R., Lu, X., Luo, Y., Wang, Z., Zhu, Y., Yin, M., Ding, Y., and Lin, J. (2020). TUSC3 induces drug resistance and cellular stemness via Hedgehog signaling pathway in colorectal cancer. Carcinogenesis 41, 1755–1766. 10.1093/carcin/bgaa038.

26. Jeon, Y.-J., Kim, T., Park, D., Nuovo, G.J., Rhee, S., Joshi, P., Lee, B.-K., Jeong, J., Suh, S., Grotzke, J.E., et al. (2018). miRNA-mediated TUSC3 deficiency enhances UPR and ERAD to promote metastatic potential of NSCLC. Nat Commun 9, 5110. 10.1038/s41467-018-07561-8.

27. Yu, X., Zhai, C., Fan, Y., Zhang, J., Liang, N., Liu, F., Cao, L., Wang, J., and Du, J. (2017). TUSC3: a novel tumour suppressor gene and its functional implications. Preprint, 10.1111/jcmm.13128 https://doi.org/10.1111/jcmm.13128.

28. Pils, D., Horak, P., Gleiss, A., Sax, C., Fabjani, G., Moebus, V.J., Zielinski, C., Reinthaller, A., Zeillinger, R., and Krainer, M. (2005). Five genes from chromosomal band 8p22 are significantly down-regulated in ovarian carcinoma. Cancer 104, 2417–2429. 10.1002/cncr.21538.

29. Slutsky, I., Abumaria, N., Wu, L.-J., Huang, C., Zhang, L., Li, B., Zhao, X., Govindarajan, A., Zhao, M.-G., Zhuo, M., et al. (2010). Enhancement of Learning and Memory by Elevating Brain Magnesium. Neuron 65, 165–177. 10.1016/j.neuron.2009.12.026.

30. Zhou, H., and Clapham, D.E. (2009). Mammalian MagT1 and TUSC3 are required for cellular magnesium uptake and vertebrate embryonic development. Proceedings of the National Academy of Sciences 106, 15750–15755. 10.1073/pnas.0908332106.

31. Garshasbi, M., Kahrizi, K., Hosseini, M., Nouri Vahid, L., Falah, M., Hemmati, S., Hu, H., Tzschach, A., Ropers, H.H., Najmabadi, H., et al. (2011). A novel nonsense mutation in TUSC3 is responsible for non-syndromic autosomal recessive mental retardation in a consanguineous Iranian family. Am J Med Genet A 155, 1976–1980. 10.1002/ajmg.a.34077.

32. Ninagawa, S., George, G., and Mori, K. (2021). Mechanisms of productive folding and endoplasmic reticulum-associated degradation of glycoproteins and non-glycoproteins. Preprint, 10.1016/j.bbagen.2020.129812 https://doi.org/10.1016/j.bbagen.2020.129812.

33. Xun, Q., Bi, C., Cui, X., Wu, H., Wang, M., Liao, Y., Wang, R., Xie, H., Shen, Z., and Fang, M. (2018). MagT1 is essential for Drosophila development through the shaping of Wingless and Decapentaplegic signaling pathways. Biochem Biophys Res Commun 503, 1148–1153. 10.1016/j.bbrc.2018.06.133.

34. Galeone, A., Han, S.Y., Huang, C., Hosomi, A., Suzuki, T., and Jafar-Nejad, H. (2017). Tissue-specific regulation of BMP signaling by Drosophila N-glycanase 1. Elife 6, e27612. 10.7554/eLife.27612.

35. Sopory, S., Kwon, S., Wehrli, M., and Christian, J.L. (2010). Regulation of Dpp activity by tissue-specific cleavage of an upstream site within the prodomain. Dev Biol 346, 102–112. 10.1016/J.YDBIO.2010.07.019.

36. Hang, Q., Zhou, Y., Hou, S., Zhang, D., Yang, X., Chen, J., Ben, Z., Cheng, C., and Shen, A. (2014). Asparagine-linked glycosylation of bone morphogenetic protein-2 is required for secretion and osteoblast differentiation. Glycobiology 24, 292–304. 10.1093/glycob/cwt110.

37. Kim, H.-S., Sanchez, M.L., Silva, J., Schubert, H.L., Dennis, R., Hill, C.P., and Christian, J.L. (2025). Mutations that prevent phosphorylation of the BMP4 prodomain impair proteolytic maturation of homodimers leading to lethality in mice. Elife 14, RP105018. 10.7554/eLife.105018.

38. Reily, C., Stewart, T.J., Renfrow, M.B., and Novak, J. (2019). Glycosylation in health and disease. Preprint, 10.1038/s41581-019-0129-4 https://doi.org/10.1038/s41581-019-0129-4.

39. Akiyama, T., Raftery, L.A., and Wharton, K.A. (2024). Bone morphogenetic protein signaling: the pathway and its regulation. Preprint, 10.1093/genetics/iyad200 https://doi.org/10.1093/genetics/iyad200.

40. Tauscher, P.M., Gui, J., and Shimmi, O. (2016). Adaptive protein divergence of BMP ligands takes place under developmental and evolutionary constraints. Development (Cambridge) 143. 10.1242/dev.130427.

41. Galeone, A., Adams, J.M., Matsuda, S., Presa, M.F., Pandey, A., Han, S.Y., Tachida, Y., Hirayama, H., Vaccari, T., Suzuki, T., et al. (2020). Regulation of BMP4/Dpp retrotranslocation and signaling by deglycosylation. Elife 9, e55596. 10.7554/eLife.55596.

42. Zi, Z., Chapnick, D.A., and Liu, X. (2012). Dynamics of TGF-β/Smad signaling. FEBS Lett 586, 1921–1928. 10.1016/j.febslet.2012.03.063.

43. Groppe, J., Rumpel, K., Economides, A.N., Stahl, N., Sebald, W., and Affolter, M. (1998). Biochemical and Biophysical Characterization of Refolded <em>Drosophila</em> DPP, a Homolog of Bone Morphogenetic Proteins 2 and 4 *. Journal of Biological Chemistry 273, 29052–29065. 10.1074/jbc.273.44.29052.

44. Nelsen, S.M., and Christian, J.L. (2009). Site-specific Cleavage of BMP4 by Furin, PC6, and PC7 *. Journal of Biological Chemistry 284, 27157–27166. 10.1074/jbc.M109.028506.

45. Korennykh, A. V, Egea, P.F., Korostelev, A.A., Finer-Moore, J., Zhang, C., Shokat, K.M., Stroud, R.M., and Walter, P. (2009). The unfolded protein response signals through high-order assembly of Ire1. Nature 457, 687–693. 10.1038/nature07661.

46. Zhang, K., and Kaufman, R.J. (2004). Signaling the Unfolded Protein Response from the Endoplasmic Reticulum *. Journal of Biological Chemistry 279, 25935–25938. 10.1074/jbc.R400008200.

47. Kim, W., Spear, E.D., and Ng, D.T.W. (2005). Yos9p Detects and Targets Misfolded Glycoproteins for ER-Associated Degradation. Mol Cell 19, 753–764. 10.1016/j.molcel.2005.08.010.

48. Alcock, F., and Swanton, E. (2009). Mammalian OS-9 Is Upregulated in Response to Endoplasmic Reticulum Stress and Facilitates Ubiquitination of Misfolded Glycoproteins. J Mol Biol 385, 1032–1042. 10.1016/j.jmb.2008.11.045.

49. Lai, C.W., Aronson, D.E., and Snapp, E.L. (2010). BiP Availability Distinguishes States of Homeostasis and Stress in the Endoplasmic Reticulum of Living Cells. Mol Biol Cell 21, 1909–1921. 10.1091/mbc.e09-12-1066.

50. Freeze, H.H., and Kranz, C. (2010). Endoglycosidase and Glycoamidase Release of N-Linked Glycans. Curr Protoc Mol Biol 89, 17.13A.1–17.13A.25. 10.1002/0471142727.mb1713as89.

51. Zhang, M., Xiao, J., Liu, J., Bai, X., Zeng, X., Zhang, Z., and Liu, F. (2023). Calreticulin as a marker and therapeutic target for cancer. Clin Exp Med 23, 1393–1404. 10.1007/s10238-022-00937-7.

52. Linstedt, A.D., and Hauri, H.P. (1993). Giantin, a novel conserved Golgi membrane protein containing a cytoplasmic domain of at least 350 kDa. Mol Biol Cell 4, 679–693. 10.1091/mbc.4.7.679.

53. Chardin, P., and McCormick, F. (1999). Brefeldin A: The Advantage of Being Uncompetitive. Cell 97, 153–155. 10.1016/S0092-8674(00)80724-2.

54. D’Alonzo, D., De Fenza, M., Porto, C., Iacono, R., Huebecker, M., Cobucci-Ponzano, B., Priestman, D.A., Platt, F., Parenti, G., Moracci, M., et al. (2017). N-Butyl-l-deoxynojirimycin (l-NBDNJ): Synthesis of an Allosteric Enhancer of α-Glucosidase Activity for the Treatment of Pompe Disease. J Med Chem 60, 9462–9469. 10.1021/acs.jmedchem.7b00646.

55. Anderson, D.J., Le Moigne, R., Djakovic, S., Kumar, B., Rice, J., Wong, S., Wang, J., Yao, B., Valle, E., Kiss von Soly, S., et al. (2015). Targeting the AAA ATPase p97 as an Approach to Treat Cancer through Disruption of Protein Homeostasis. Cancer Cell 28. 10.1016/j.ccell.2015.10.002.

56. Ballar, P., Pabuccuoglu, A., and Kose, F.A. (2011). Different p97/VCP complexes function in retrotranslocation step of mammalian Er-associated degradation (ERAD). International Journal of Biochemistry and Cell Biology 43. 10.1016/j.biocel.2010.12.021.

57. Chen, Y., Hu, D., Yabe, R., Tateno, H., Qin, S.-Y., Matsumoto, N., Hirabayashi, J., and Yamamoto, K. (2011). Role of malectin in Glc2Man9GlcNAc2-dependent quality control of α1-antitrypsin. Mol Biol Cell 22, 3559–3570. 10.1091/mbc.e11-03-0201.

58. Wu, Y., Li, Q., and Chen, X.-Z. (2007). Detecting protein–protein interactions by far western blotting. Nat Protoc 2, 3278–3284. 10.1038/nprot.2007.459.

59. Song, J. (2013). Why do proteins aggregate? “Intrinsically insoluble proteins”? And “dark mediators”? Revealed by studies on “insoluble proteins”? Solubilized in pure water. F1000Res 2. 10.12688/f1000research.2-94.v1.

60. Padgett, R.W., St. Johnston, R.D., and Gelbart, W.M. (1987). A transcript from a Drosophila pattern gene predicts a protein homologous to the transforming growth factor-β family. Nature 325. 10.1038/325081a0.

61. Nakayama*, T., Cui*, Y., and Christian**, J.L. (2000). Regulation of BMP/Dpp signaling during embryonic development. Cell Mol Life Sci 57, 943–956. 10.1007/PL00000736.

62. Hursh, D.A., Padgett, R.W., and Gelbart, W.M. (1993). Cross regulation of decapentaplegic and Ultrabithorax transcription in the embryonic visceral mesoderm of Drosophila. Development 117, 1211–1222. 10.1242/dev.117.4.1211.

63. Panganiban, G.E.F., Reuter, R., Scott, M.P., and Hoffmann, F.M. (1990). A Drosophila growth factor homolog, decapentaplegic, regulates homeotic gene expression within and across germ layers during midgut morphogenesis. Development 110, 1041–1050. 10.1242/dev.110.4.1041.

64. Li, H., Qi, Y., and Jasper, H. (2016). Ubx dynamically regulates Dpp signaling by repressing Dad expression during copper cell regeneration in the adult Drosophila midgut. Dev Biol 419, 373–381. 10.1016/j.ydbio.2016.08.027.

65. Newfeld, S.J., Chartoff, E.H., Graff, J.M., Melton, D.A., and Gelbart, W.M. (1996). Mothers against dpp encodes a conserved cytoplasmic protein required in DPP/TGF-β responsive cells. Development 122, 2099–2108. 10.1242/dev.122.7.2099.

66. Akiyama, T., and Gibson, M.C. (2015). Decapentaplegic and growth control in the developing Drosophila wing. Nature 527, 375–378. 10.1038/nature15730.

67. Bischof, J., Maeda, R.K., Hediger, M., Karch, F., and Basler, K. (2007). An optimized transgenesis system for Drosophila using germ-line-specific φC31 integrases. Proceedings of the National Academy of Sciences 104, 3312–3317. 10.1073/pnas.0611511104.

68. Venken, K.J.T., He, Y., Hoskins, R.A., and Bellen, H.J. (2006). P[acman]: A BAC Transgenic Platform for Targeted Insertion of Large DNA Fragments in D. melanogaster. Science (1979) 314, 1747–1751. 10.1126/science.1134426.

69. Xu, L., Liu, X., Peng, F., Zhang, W., Zheng, L., Ding, Y., Gu, T., Lv, K., Wang, J., Ortinau, L., et al. (2020). Protein quality control through endoplasmic reticulum-associated degradation maintains haematopoietic stem cell identity and niche interactions. Nat Cell Biol 22. 10.1038/s41556-020-00581-x.

70. Shenkman, M., Ogen-Shtern, N., Patel, C., Groisman, B., Pasmanik-Chor, M., Schermann, S.M., Körner, R., and Lederkremer, G.Z. (2025). Oligosaccharyltransferase is involved in targeting to ER-associated degradation. bioRxiv, 2024.05.12.593735. 10.1101/2024.05.12.593735.

71. Garshasbi, M., Hadavi, V., Habibi, H., Kahrizi, K., Kariminejad, R., Behjati, F., Tzschach, A., Najmabadi, H., Ropers, H.H., and Kuss, A.W. (2008). A Defect in the TUSC3 Gene Is Associated with Autosomal Recessive Mental Retardation. Am J Hum Genet 82. 10.1016/j.ajhg.2008.03.018.

72. Molinari, F., Foulquier, F., Tarpey, P.S., Morelle, W., Boissel, S., Teague, J., Edkins, S., Futreal, P.A., Stratton, M.R., Turner, G., et al. (2008). Oligosaccharyltransferase-Subunit Mutations in Nonsyndromic Mental Retardation. Am J Hum Genet 82. 10.1016/j.ajhg.2008.03.021.

73. El Chehadeh, S., Bonnet, C., Callier, P., Béri, M., Dupré, T., Payet, M., Ragon, C., Mosca-Boidron, A.L., Marle, N., Mugneret, F., et al. (2015). Homozygous truncating intragenic duplication in TUSC3 responsible for rare autosomal recessive nonsyndromic intellectual disability with no clinical or biochemical metabolic markers. In JIMD Reports 10.1007/8904_2014_390.

74. Ramírez, A.S., Kowal, J., and Locher, K.P. (2019). Cryo–electron microscopy structures of human oligosaccharyltransferase complexes OST-A and OST-B. Science (1979) 366, 1372–1375. 10.1126/science.aaz3505.

