## Supplementary material for "TUSC3 serves as a rate-limiting gatekeeper of a glycan-mediated ER Triage Checkpoint for BMP4/Dpp": Figure supplement 1-3

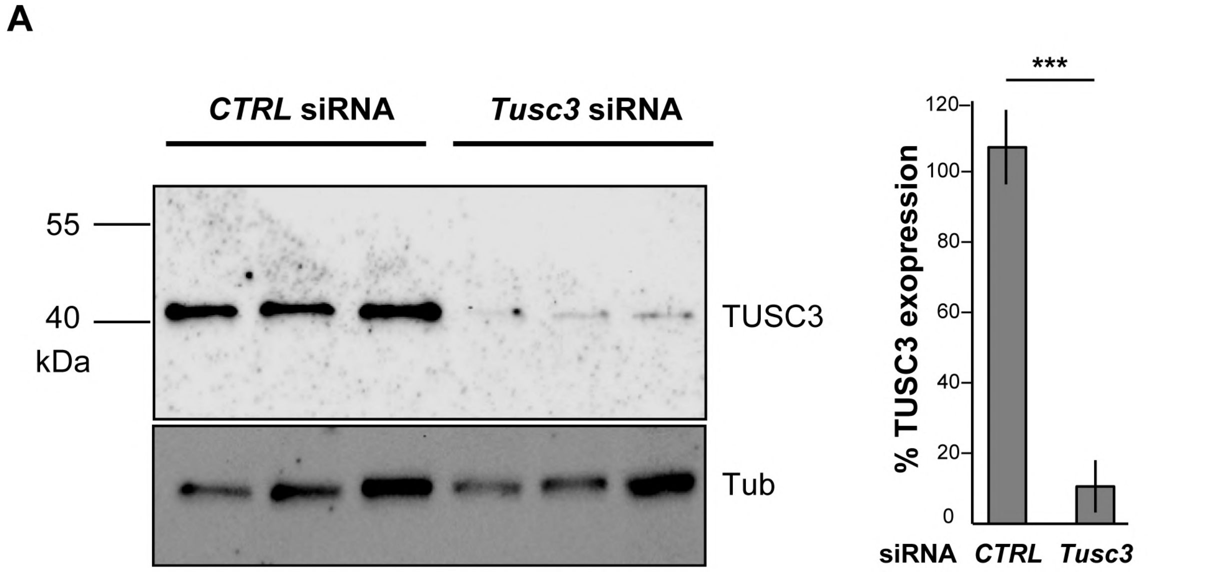


**Figure S1.** ***Tusc3*** **is efficiently silenced in MEFs by siRNA.** (A) Immunoblot analysis of TUSC3 and Tubulin in MEFs transfected with control (*CTRL*) and *Tusc3* siRNA 48 h post-transfection. Quantification (right) shows TUSC3 protein levels normalized to Tubulin (mean ± SEM, n = 3; ***p < 0.001, Student's t test).

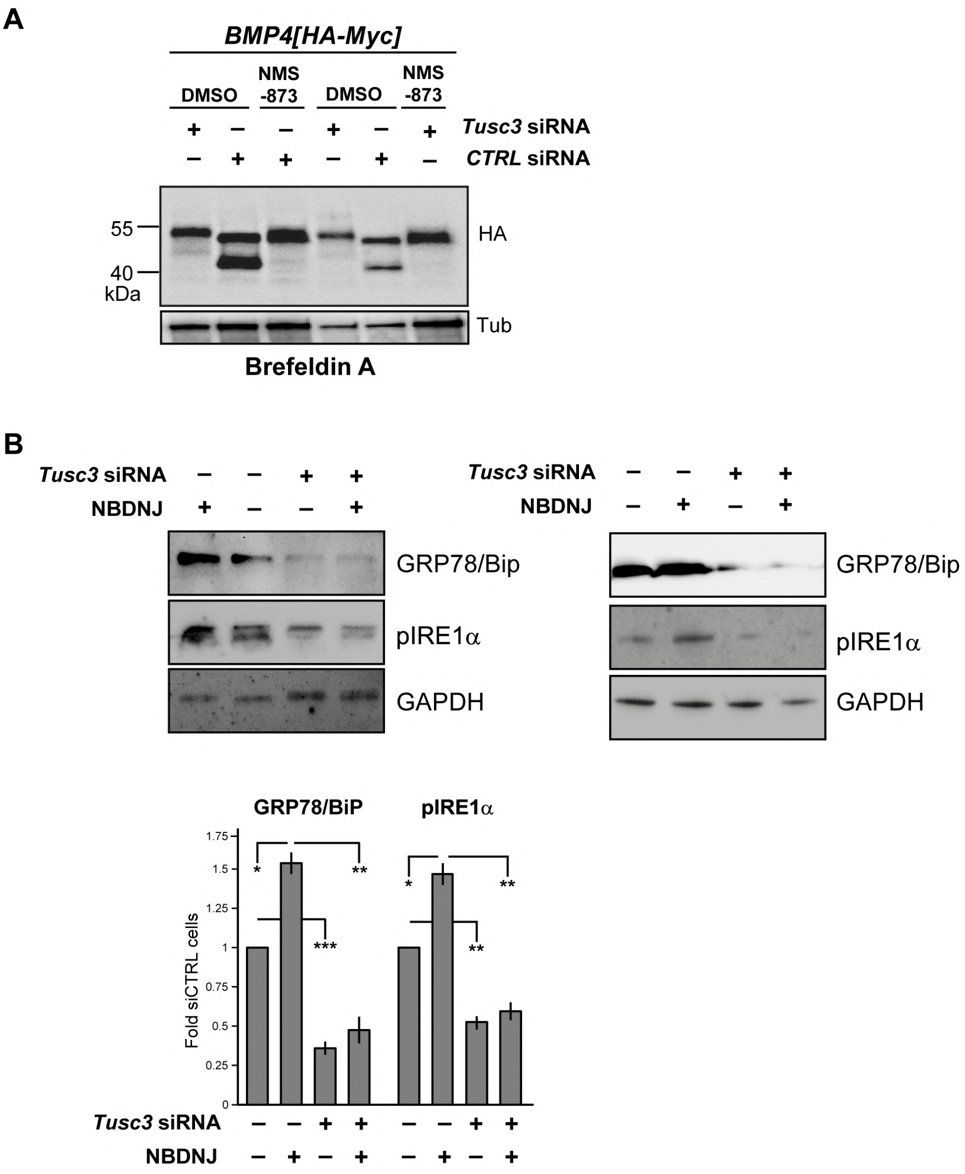


**Figure S2.** ***Tusc3*** **knockdown alters BMP4 glycoform processing and ER stress responses.** (A) Western blot of BMP4-HA in *Tusc3*-knockdown MEFs treated with the VCP inhibitor NMS-873 or with DMSO as control. (B) Western blots with the indicated antibodies on extracts from MEFs treated with *Tusc3* siRNA and or the GCS inhibitor NBDNJ. Graph shows the quantification of BiP and pIRE1α expression levels in control, *Tusc3*-knockdown, and NBDNJ-treated MEFs. Data normalized to GAPDH. n = 3 biological replicates.

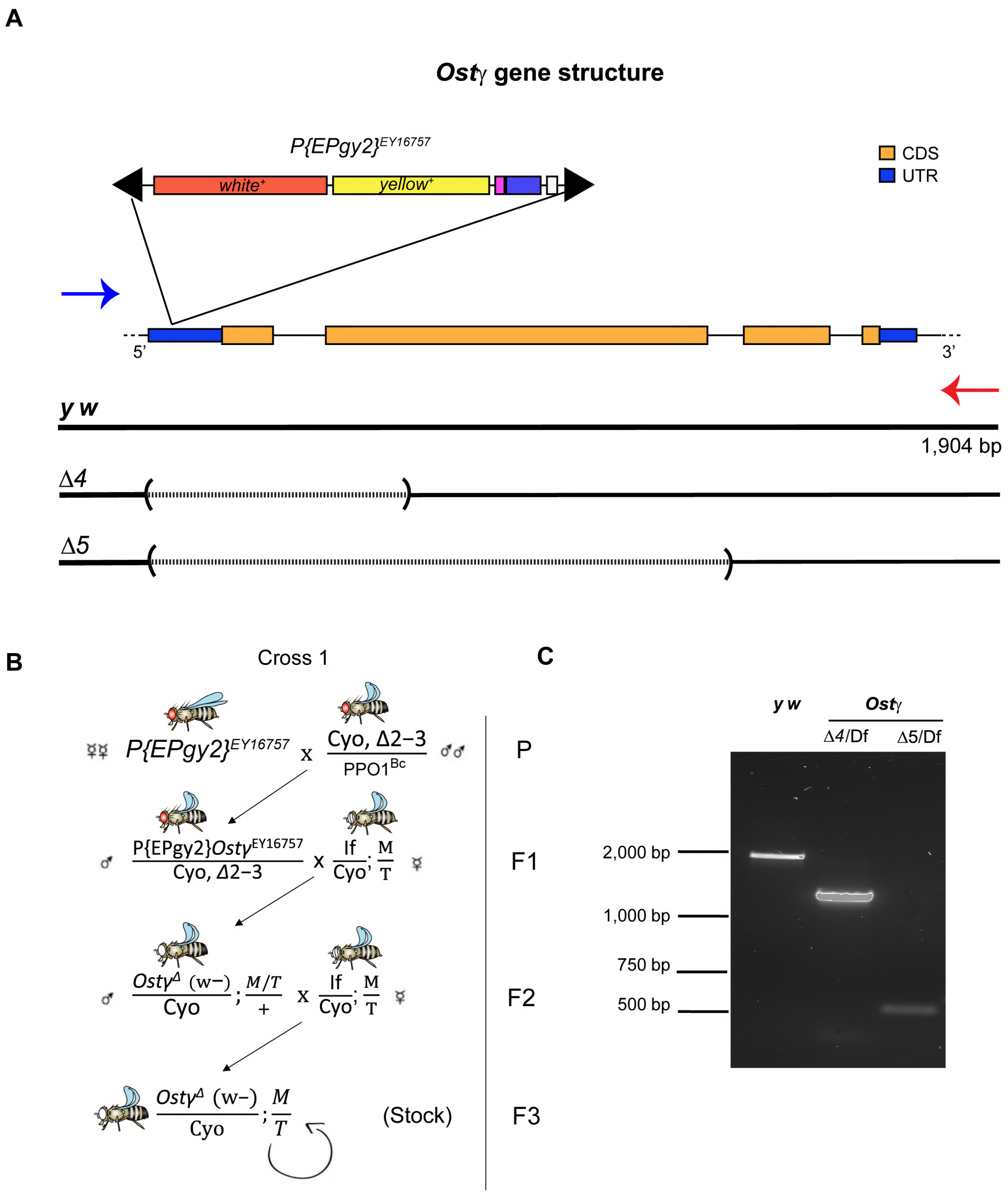


**Figure S3. Molecular and phenotypic validation of *Ostγ* alleles in *Drosophila*.** (A) Schematic of the *Ostγ* locus on chromosome 2L showing the *P{EPgy2}* insertion site in the *5' UTR* (*EY16757*). Blue and red arrows mark the location of the PCR primers used in panel C. Dashed lines indicate the deleted region in each allele. (B) Schematic of genetic crosses designed to excise the *P{EPgy2}* insertion using Δ2-3 transposase. F3 males were balanced and stocked for PCR-based screening at the *Ostγ* locus. (C) PCR-based amplification of the *Ostγ* locus using primers flanking the *Ostγ* gene from genomic DNA the indicated genotype.
